# Pregistered movie-fMRI analyses reveal altered visual feature encoding in autism in pSTS

**DOI:** 10.64898/2026.03.23.713749

**Authors:** Jeff Mentch, Yibei Chen, Tamara Vanderwal, Satrajit S Ghosh

**Affiliations:** Program in Speech and Hearing Bioscience and Technology, Harvard University, Cambridge, United States; McGovern Institute for Brain Research, Massachusetts Institute of Technology, Cambridge, United States; Department of Psychiatry, University of British Columbia, Vancouver, British Columbia, Canada; BC Children’s Hospital Research Institute, Vancouver, British Columbia, Canada; Department of Otolaryngology, Harvard Medical School, Boston, United States

## Abstract

Sensory–perceptual differences are widely reported in autism, yet their underlying mechanisms remain unclear. We tested preregistered hypotheses using stacked encoding models applied to naturalistic movie-viewing fMRI from children and adolescents with and without an autism diagnosis from the Healthy Brain Network. We mapped cortical responsiveness to low- and high-level auditory and visual feature spaces. Contrary to enhanced perceptual functioning predictions, autism was not associated with increased low-level encoding in primary sensory cortices. Instead, autistic children and adolescents had reduced high-level visual representations and a relative shift toward low-level over high-level feature encoding in integration and social brain regions including the pSTS and adjacent face/social areas. In pSTS, this high–low weighting tracked Social Responsiveness Scale (SRS) scores. By contrast, audio–visual modality preference and sensory dominance were broadly conserved across groups. Developmentally, encoding exhibited strong, lateralized, modality-congruent age effects. Together, these findings favor weak central coherence accounts over early sensory enhancement, constrain mechanisms to altered visual feature weighting within social/multisensory networks, and demonstrate the value of naturalistic stimuli and encoding models for characterizing sensory-perceptual neurodevelopmental differences.

## Introduction

Sensory processing differences are a common and clinically meaningful feature of autism with a profound impact on daily life (***Ben-Sasson et al., 2007***; ***Chamak et al., 2008***). As many as 90% of autistic individuals exhibit altered sensory function (***Leekam et al., 2007***; ***Tavassoli et al., 2014***) with differences reported across all senses (***Robertson and Baron-Cohen, 2017***; ***Dakin and Frith, 2005***; ***Nieto Del Rincón, 2008***; ***Bennetto et al., 2007***; ***Balasco et al., 2020***; ***DuBois et al., 2016***), including features such as hyper- and hypo-sensitivities, enhanced perceptual abilities (***Lefebvre et al., 2023***; ***Mottron et al., 2006***; ***O’riordan, 2004***; ***Mottron and Burack, 2001***), altered susceptibility to perceptual illusions (***Happé, 1996***; ***Feldman et al., 2022***), and differences in multisensory integration (MSI) (***Stevenson et al., 2014b***,a; ***Feldman et al., 2018***). These traits emerge early, track with social and cognitive outcomes, and vary across individuals and modalities (***Uljarević et al., 2017***; ***Williams et al., 2023***; ***Georgiades et al., 2017***; ***Chen et al., 2022***; ***Simonoff et al., 2020***; ***Smith et al., 2012***). Despite their prevalence and impact, the neurobiological substrates of these altered sensory functions remain relatively uncharacterized (***Marco et al., 2011***; ***Robertson and Baron-Cohen, 2017***).

### Theories of sensory differences

A number of different theories exist surrounding autistic sensory differences. Enhanced perceptual functioning (EPF) emphasizes heightened low-level processing and improved discrimination (***Mottron and Burack, 2001***; ***Mottron et al., 2006***), while weak central coherence (WCC) highlights a local-over-global perceptual preference (***Frith, 1989***; ***Happé et al., 2006***) (see ***Figure 1*** A for an illustrated comparison). Related Bayesian accounts propose altered use or updating of perceptual priors or an atypical balance between statistical likelihood and priors (***Pellicano and Burr, 2012***; ***Schneebeli et al., 2022***; ***Brock, 2012***) and further, predictive-coding, a process-level instantiation of these Bayesian ideas, implicates imbalances between prediction errors and top-down predictions (***Mumford, 1992***; ***Rao and Ballard, 1999***; ***Friston, 2005***; ***Lawson et al., 2014***; ***Van de Cruys et al., 2014***; ***Lawson et al., 2017***). More mechanistically, neurobiological accounts emphasize excitation–inhibition (E/I) imbalances and neural noise, with support from EEG, genetics, and animal studies (***Rubenstein and Merzenich, 2003***; ***Dinstein et al., 2012***; ***Puts et al., 2017***; ***Wei et al., 2021***; ***Hollestein et al., 2023***; ***Han et al., 2012***; ***Robertson et al., 2016***). An integrative view links E/I alterations to disrupted divisive normalization central to causal inference and predictive coding (***Noel and Angelaki, 2023***). In recent years, “Sensory-first” accounts of autism argue that perceptual differences precede and scaffold downstream social–communicative phenotypes (***Levit-Binnun et al., 2013***; ***Cascio et al., 2016***; ***Robertson and Baron-Cohen, 2017***; ***Falck-Ytter and Bussu, 2023***; ***Russo et al., 2025***; ***Takesian and Hensch, 2013***).

**Figure 1.**
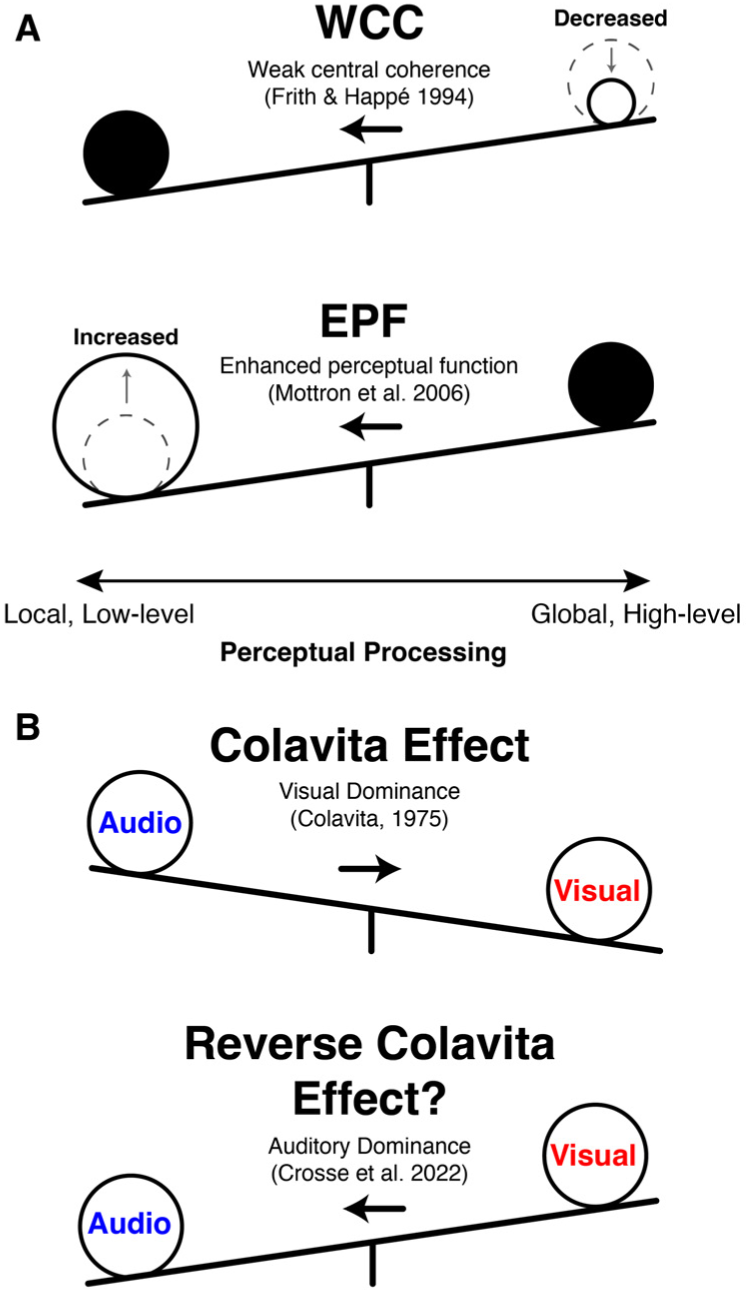
(A) A visual depiction contrasting how perceptual processing is thought to be different in autism across the two prominent theories of WCC and EPF. (B) An illustration comparing visual sensory dominance (the Colavita effect) with decreased visual dominance or auditory sensory dominance (a reverse Colavita effect), which has been observed in autism.

Overall, these varied accounts yield some testable predictions for naturalistic representations, among them: relatively amplified low-level and reduced high-level encoding and altered local–global and unimodal–multisensory balance;

Another axis of sensory organization relevant to autism is the balance between auditory and visual channels during naturalistic perception. Classic multisensory work shows that audiovisual (AV) integration engages regions including the superior temporal sulcus (STS), which can be supra-additive for congruent AV events (***Calvert et al., 2000***, ***2001***), with multisensory influences present even in regions often treated as unisensory (***Ghazanfar and Schroeder, 2006***; ***Stein and Stanford, 2008***). Developmentally, AV integration is protracted, approaching adult-like levels around late childhood, and may follow altered trajectories in autism (***Neil et al., 2006***; ***Burr and Gori, 2012***; ***Russo et al., 2025***). Behaviorally, autism has been associated with differences in multisensory integration and temporal binding (e.g., widened temporal binding windows and reduced MSI enhancements), with downstream implications for domains including speech processing (***Stevenson et al., 2014b***,a; ***Feldman et al., 2018***; ***Wang et al., 2024***). Classic work on sensory dominance shows a robust visual-dominance in adults (the Colavita effect) that develops from earlier auditory predominance in childhood (***Colavita, 1974***; ***Hirst et al., 2018***; ***Ross et al., 2021***). In autism, findings include reduced or reversed visual dominance, including a reverse Colavita effect and a shift toward auditory bias (***Ross et al., 2021***; ***Moro et al., 2012***; ***O’connor and Hermelin, 1965***), hinting at potentially altered audio vs. visual sensory feature weightings in autism (see ***Figure 1*** B).

### Developmental maturation of sensory and association systems

Beyond autism, it is useful to note several robust developmental trends. Primary sensory cortices mature relatively early, whereas higher-order association regions (e.g., pSTS, TPJ, FFA) show protracted structural–functional specialization into adolescence (***Thompson and Nelson, 2001***; ***Sydnor et al., 2021***). Multisensory processing also sharpens with age: audiovisual temporal binding windows narrow, and cross-modal influences in STS become more reliable and congruent with task demands (***Neil et al., 2006***; ***Burr and Gori, 2012***). These maturational trends imply that group differences observed in association cortices and multisensory networks must be interpreted against a baseline of ongoing developmental tuning.

### Neuroimaging evidence for altered sensory processing in autism

Within autism, neuroimaging findings have consistently implicated primary sensory brain regions (e.g. motion-related differences in visual cortex (***Robertson et al., 2014***), enlarged population receptive fields (***Schwarzkopf et al., 2014***), and heightened connectivity within primary visual and auditory cortices (***Martínez et al., 2020***; ***Cerliani et al., 2015***)). Increased trial-to-trial variability of evoked responses has also been observed across early visual, auditory, and somatosensory cortices (***Dinstein et al., 2012***). Atypical early auditory discrimination has been further measured through mismatch negativity (MMN), with MMN amplitudes directly linked to sensory over-responsivity and autistic traits (***Cary et al., 2024***). During audiovisual integration tasks, autistic participants were found to exhibit auditory-led phase resets in visual cortex, opposite the visual-led synchronization typical of neurotypical observers, consistent with altered precision and reliability weighting of sensory cues and tying primary sensory processing alterations to audiovisual perception (***Ronconi et al., 2023***).

Alongside these early sensory processing differences, a growing body of research highlights significant involvement of higher-order cortical regions. These include but are not limited to the temporoparietal junction (TPJ; found to have decreased cortical thickness (***You et al., 2024***) and altered connectivity to the cerebellum (***Igelström et al., 2017***)), the posterior superior temporal sulcus (pSTS; found to have reduced activation during biological motion perception (***Koldewyn et al., 2011***; ***Federici et al., 2020***) and altered intrinsic and dynamic functional connectivity (***Alaerts et al., 2015***, ***2014***; ***Shih et al., 2011***; ***Guo et al., 2019***)), and the fusiform face area (FFA; found to have altered volume and hemispheric asymmetry (***Herbert et al., 2002***; ***Floris et al., 2025***) along with reduced activation in response to faces (***Pierce et al., 2001***; ***Schultz et al., 2003***; ***Hadjikhani et al., 2007***), though some later studies found no difference for familiar faces (***Pierce et al., 2004***; ***Hadjikhani et al., 2007***)).

Beyond specific low- and high-level regions with altered connectivity, there have been widespread reports of more generally altered connectivity across the brain, in the form of under-connectivity (***Damarla et al., 2010***; ***Abrams et al., 2013***; ***Alaerts et al., 2014***), over-connectivity (***Rudie and Dapretto, 2013***; ***Keown et al., 2013***), and differing local and global connectivity (***Courchesne and Pierce, 2005***). These patterns are thought to be dynamic and changing with age (***Lawrence et al., 2019***). Some findings point to an anterior to posterior gradient across the brain of under- to over-connectivity in autism, perhaps explaining enhanced perceptual performance with increased local connectivity in early sensory areas and disrupted social behavior with decreased local connectivity in frontal areas (***Keown et al., 2013***; ***Supekar et al., 2013***; ***Di Martino et al., 2014***; ***Rudie and Dapretto, 2013***). While most studies assess activation or connectivity, fewer examine what or how information is represented during rich, dynamic perception and a clear gap remains.

### Naturalistic stimuli and autism

Naturalistic stimuli (such as movies and narratives) in neuroimaging studies can improve ecological validity and compliance while also implicitly including rich sensory, semantic, and social features (***Sonkusare et al., 2019***; ***Nastase et al., 2020***; ***Vanderwal et al., 2019***; ***Redcay and Moraczewski, 2020***; ***Haxby et al., 2014***). In autism, movie paradigms have revealed reduced ISC and ISFC and idiosyncratic responses (foundational and follow-up work by ***Hasson et al. 2009***; ***Salmi et al. 2013***;***Byrge et al. 2015***), links to comprehension and symptomatology (e.g., default-mode/control networks; Bolton and colleagues (***2018***; ***2020***)), region- or event-specific effects (TPJ, theory-of-mind; ***Pantelis et al. 2015***; ***Lyons et al. 2020***), and cross-national replication of ISFC reductions (***Lin et al., 2025***). Deficits in audiovisual synchrony relevant to speech have also been reported (***Wang et al., 2024***). Most work to date leverages metrics of brain synchrony, while feature-resolved accounts of what and where information may be differentially represented in these naturalistic contexts remains relatively unexplored.

### Encoding models

Encoding models enable the direct mapping of stimulus features to brain responses, allowing us to quantify the representation of information across the brain (***Naselaris et al., 2011***; ***Diedrichsen and Kriegeskorte, 2017***). Encoding models are readily adaptable to high-dimensional, naturalistic features (***Samara et al., 2025***; ***Khosla et al., 2021***; ***Huth et al., 2016***; ***Haxby et al., 2011***) and some specific implementations, such as as stacked encoding models (***Lin et al., 2024***) and banded regression (***Dupré la Tour et al., 2022***) enable joint tests across feature spaces. In stacked encoding models, instead of relying solely on the predictive performance of separate feature models, the secondary “stacking” stage yields weights that index the relative contribution of each feature space to the joint prediction, providing, for example, a direct measure of high- vs. low-level or audio vs. visual perceptual preferences when these feature classes compete to explain the same responses. Despite their successful applications in vision and language research, encoding model approaches are relatively underutilized in autism research relative to ISC based methods, particularly in studies using naturalistic designs (***Dupré la Tour et al., 2025***; ***Kriegeskorte and Douglas, 2019***).

### This study: dataset, features, analyses, preregistration, hypotheses

Here, naturalistic movie-viewing fMRI data from children and adolescents with and without autism is analyzed to test a set of preregistered hypotheses (accessible at osf.io/h92gr and osf.io/47kj6; summarized in ***Table 1***). Feature sets are constructed spanning low-level (e.g., brightness, motion and loudness) through high-level (faces, bodies, and audio categories such as speech and music), separately for audio and visual streams and feed these into whole-brain grayordinate-level encoding models that quantify the explained variance for each feature class. Primary hypotheses: (i) Relative to non-autistic peers, autistic participants will show greater encoding of *low-level* perceptual features in primary sensory and other perceptual regions (H1.1), with a shift in stacked-weight preference toward low- over high-level features (H1.2); (ii) autistic auditory and visual cortices will be more unimodal (H2.1), preferentially encoding modality-congruent information and exhibiting reduced visual dominance (H2.2). Secondary hypotheses: Encoding-model metrics will covary with clinical phenotype including autism severity and sensory symptoms (H1.3, H2.3).

**Table 1.** Table of preregistered hypotheses (accessible at osf.io/h92gr and osf.io/47kj6).

|  | Hypothesis | Feature sets / metrics |
| --- | --- | --- |
| **Low- vs. high-level encoding** |  |  |
| 1.1 | Primary perceptual cortices will encode more information about low-level perceptual features in an autistic relative to a non-autistic group. | Low-level audio and visual features; $R^2$, $R_u^2$ |
| 1.2 | Perceptual regions of the brain will be more tuned towards low- as opposed to high-level sensory information in an autistic relative to a non-autistic group. | Low- vs. high- level audio and visual feature classes; $(W_{HA}-W_{LA})$, $(W_{HV}-W_{LV})$ |
| 1.3 | Encoding model-derived differences will be related to autism severity and sensory symptoms. | SRS/SSS associations with high- and low-level, audio and visual $R^2$, $R_u^2$, $(W_H-W_L)$ |
| **Audio vs. visual encoding** |  |  |
| 2.1 | Auditory and visual regions of the brain will be more *unimodal* in autistic compared to non-autistic people and encode more congruent unimodal information relative to incongruent cross-modal information. | Audio-only vs. visual-only contributions; audio and visual $R^2$, $R_u^2$ |
| 2.2 | Autistic individuals will have reduced visual dominance compared to non-autistic individuals. | Visual vs. audio stacked weights; $(W_V-W_A)$ |
| 2.3 | Audiovisual encoding model-derived differences in autistic participants will be significantly correlated with autism severity. | SRS/SSS associations with audio and visual $R^2$, $R_u^2$ and $(W_V-W_A)$ |

### Key findings

During naturalistic viewing fMRI, autistic children and adolescents have a relative shift toward low-level over high-level feature encoding in integration and social brain regions (including the pSTS), with a broadly conserved audio–visual feature balance but robust age-related effects. These findings inform sensory theories, argue against large, pervasive shifts in modality dominance, and demonstrate the utility of naturalistic encoding model methods for the investigation of sensory-perceptual neurodevelopmental differences.

## Results

### Conventions

We report explained variance (*R*^2^), unique explained variance (*R_u_*^2^, from variance partitioning), and stacked encoding model weights (*W*) for audio and visual models. Perceptual preference indices are defined as *W_V_* − *W_A_* (visual vs. audio modality preference) and *W_H_* − *W_L_* (high- vs. low-level feature preference), with positive values indicating visual (or high-level) preference and negative values indicating audio (or low-level) preference. Unless otherwise noted, all p-values are FDR-corrected across parcels, and results are shown across three framewise displacement (FD) thresholds of 40%, 60%, and 80%.

### Encoding models recover known sensory hierarchies

Aggregating across all (both autistic and non-autistic) participants, audiovisual stacked encoding models produced patterns of performance and feature representation in audio and visual cortical regions that are in keeping with known sensory hierarchies. This is evident by examining the perceptual preference index (the difference in visual and auditory model stacked weights (*W_V_* − *W_A_*)), across both perceptual ROIs and the cortical surface more broadly (***Figure 2***). Mean *R*^2^ peaked along the superior temporal sulcus (STS), and split–half, Spearman–Brown–corrected noise ceilings (an upper bound on explainable variance given response reliability/measurement noise) were highest in parcels along the STS and in parietal/occipital cortices (***Figure 2—figure Supplement 1***). Looking at model performance by sensory modality, auditory cortices showed greater audio *R*^2^ and increased audio model weights, whereas visual cortices showed the converse pattern for visual performance and weights (***Figure 2—figure Supplement 2***). Comparing stacked low- and high-level models within audio and visual models further reproduced canonical hierarchical gradients. For vision, low-level features dominated early visual cortex (e.g., V1/V2), with relatively stronger high-level contributions in downstream ventral regions (e.g., FFC, ***Figure 2—figure Supplement 3*** A). For the audition, high-level model *R*^2^ and weights increased from A1 into association areas (A4/A5, ***Figure 2—figure Supplement 3*** B). Model weights followed this same pattern along with the high vs. low preference index (*W_H_* − *W_L_*), which was negative in early sensory cortex and approached or exceeded zero in higher-order regions (***Figure 2—figure Supplement 3*** C-F, see ***Figure 2—figure Supplement 4*** for a whole-brain mean visualization). These results establish that the feature spaces and model architecture employed here are capable of recovering known sensory hierarchies.

**Figure 2.**
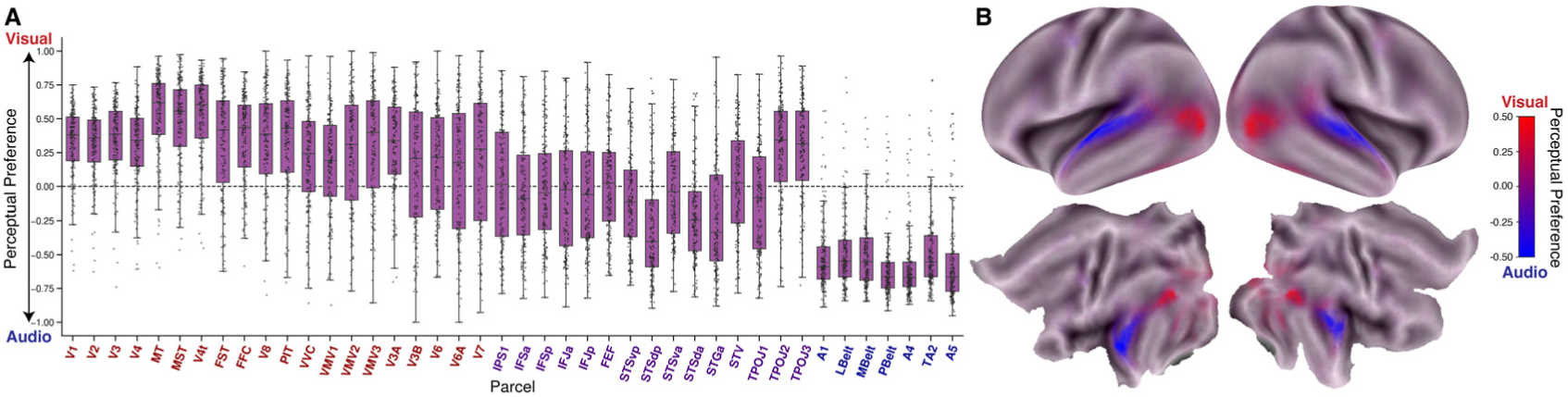
(A) Visual vs audio perceptual preference (*W_V_* − *W_A_*) for all participants (ASD and nonASD) across visual (red), audiovisual (purple), and auditory (blue) ROIs. Perceptual preference is calculated by taking the difference of visual and audio stacked model weights. Perceptual preference is in the range -1 to 1 as stacked encoding model weights range from 0 to 1. (B) Whole-brain plot of mean visual vs audio perceptual preference (*W_V_* − *W_A_*) across all participants (ASD and nonASD). This metric is the same as in A, but now plotted at each grayordinate and colored on a scale of visual (red) to audio (blue) with purple indicating regions with a relative balance between visual and audio feature weights.

### H1.1: No evidence for increased low-level encoding in primary sensory cortices in autism

The preregistered hypothesis that primary perceptual cortices contain more information about low-level perceptual features in autism was not supported. For the low-level auditory model, no significant group differences in *R*^2^ or *R_u_*^2^ were observed in primary auditory regions (all FDR q>0.05, ***Figure 3***, ***Figure 3—figure Supplement 1***). Outside auditory ROIs, some uncorrected effects appeared for *R_u_*^2^ (V4t, MST) and *R*^2^ (V4t, V7), but none survived FDR correction. Likewise, no significant group-level differences in low-level visual performance were observed in primary visual regions (all FDR q>0.05, ***Figure 3***, ***Figure 3—figure Supplement 2***). Trends in STSvp, STSdp, and STGa (uncorrected *p < .*05) did not survive FDR correciton for *R*^2^, whereas STSvp showed a significant difference for *R_u_*^2^ (40% FD; replicated at 80% FD; ***Figure 3***). Overall, early sensory-cortex encoding increases for low-level features in autism were not observed.

**Figure 3.**
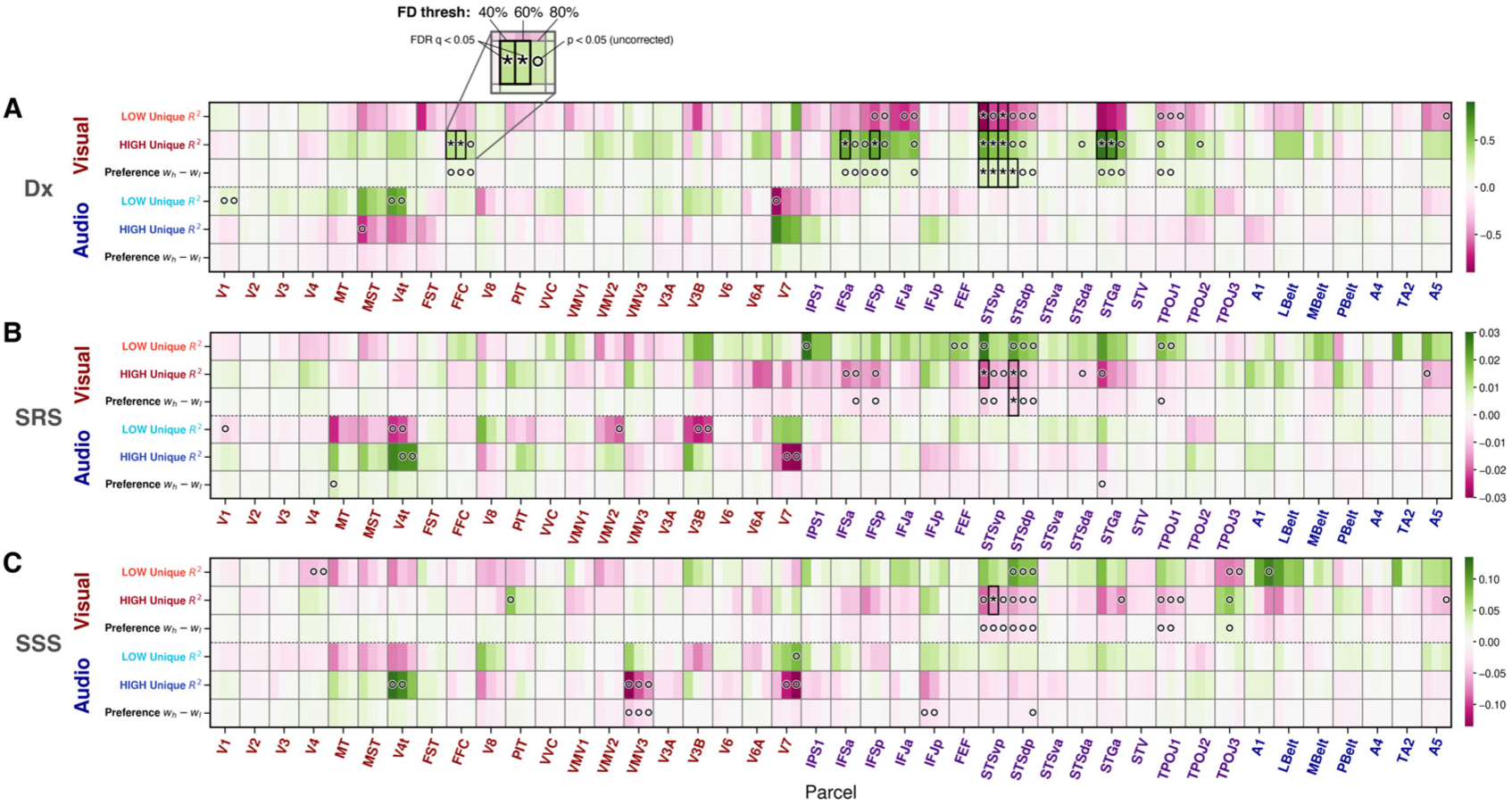
Heatmaps of fixed-effect coefficients for (A) diagnostic group (Dx: nonASD vs. ASD), (B) SRS and (C) sensory subset score (SSS), across 40%, 60% and 80% FD thresholds (left to right within each cortical parcel column). Within each of the three horizontal panels, rows denote encoding model-derived metrics (low-level *R_u_*^2^, high-level *R_u_*^2^, and their preference index (*W_H_* − *W_L_*) and columns denote Glasser ROIs ordered from early visual areas through association cortex and then auditory areas. Color indicates the magnitude and sign of the coefficient (purple=negative effect with ASD>nonASD; green=positive effect with ASD<nonASD). Asterisks (∗) mark FDR-corrected significance at q < .05; open circles (◦) mark uncorrected p < .05. The visual modality encoding models are coded with red text and the auditory with blue. Low-level visual *R_u_*^2^ was significantly different across diagnosis in STSvp and low-level audio *R_u_*^2^ was not significant anywhere (A; contradicting H1.1). Visual though not audio *W_H_* − *W_L_* was significantly different in STSvp and STSdp (A; supporting H1.2). Visual but not audio *W_H_* − *W_L_* was significantly related to SRS in STSdp at the 40% threshold (B; H1.3) but not SSS (C). Generally, results were often consistent across the three FD thresholds.

### H1.2: In autism, pSTS is weighted towards low-level visual features

Although audio models had no significant difference in low- vs. high-level preference, stacked visual models revealed a group difference in high- vs. low-level preference localized to the pSTS. Visual *W_H_* − *W_L_* was significantly lower in the autistic group in STSvp across all three FD thresholds and in STSdp at the strictest 40% threshold (***Figure 3***, ***Figure 4***).

**Figure 4.**
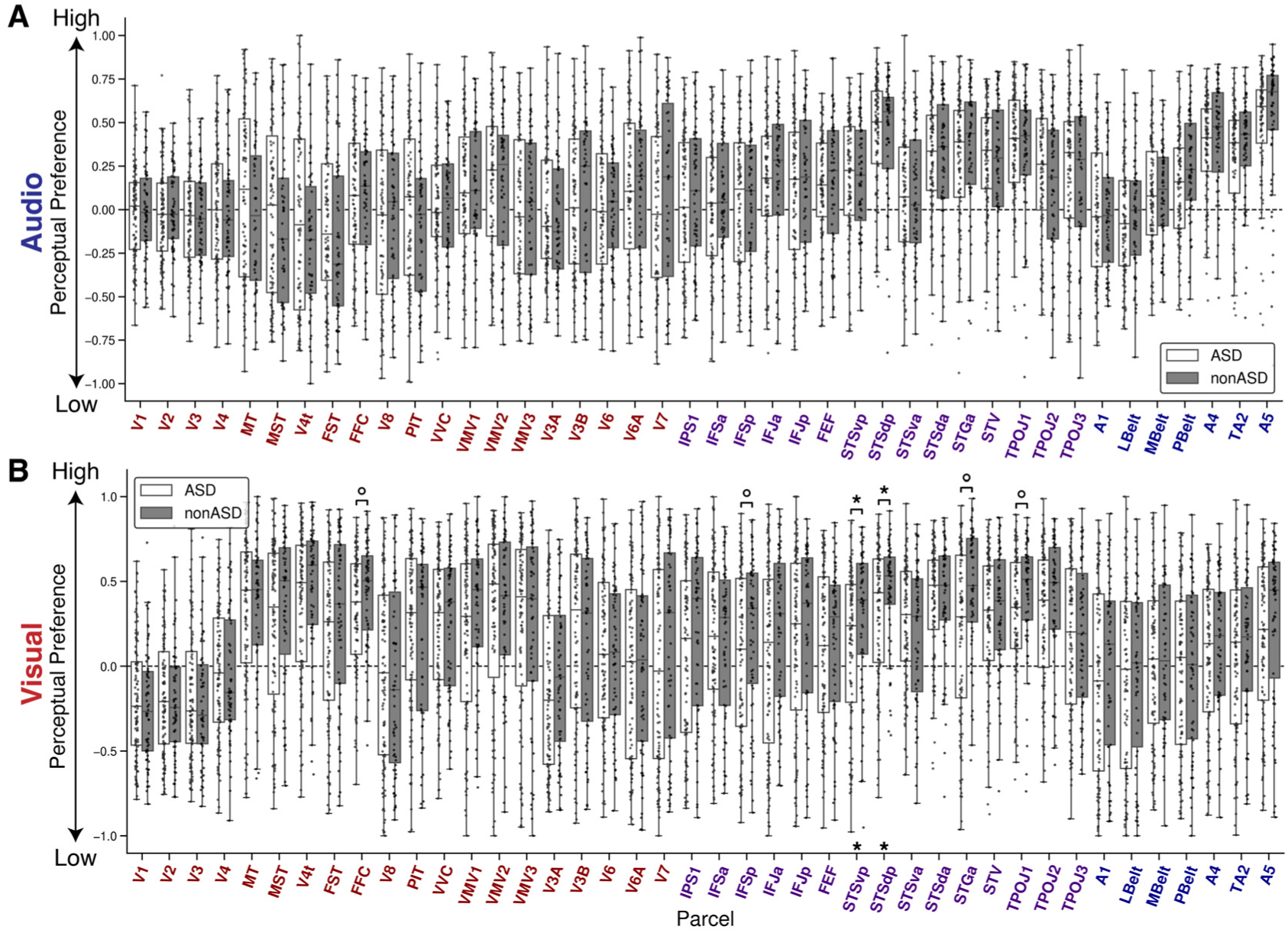
(A) Box plot of audio stacked encoding model perceptual preference across ASD (white) and nonASD (gray) groups for all perceptual ROIs. (B) Corresponding box plot of the visual stacked encoding models. Results correspond to the 40% FD threshold. Boxes annotated with an asterisk indicate a significant group difference (FDR q<0.05) while a circle indicates an initially significant difference between groups that did not survive FDR correction. Boxes show the quartiles of the dataset and whiskers show the distribution with the exception of outliers. Each dot is the mean perceptual preference from a single subject from all statistically significant grayordinates within each ROI.

### H1.3: pSTS encoding differences are related to autism severity as measured by the SRS

Across visual and auditory models and the SRS and SSS measures, visual high- vs. low-level preference was significantly related to SRS after FDR correction in STSdp at the 40% threshold where lower *W_H_* − *W_L_* (increased low-level preference) was associated with higher SRS score (coefficients summarized in ***Figure 3***).

### Exploratory: High-level visual representations are reduced in face and social brain regions in autism

Although the preregistered hypotheses focused on low-level feature representation and the balance of low- and high-level feature weights, the stacked models also provide high-level feature representation metrics. High-level audio *R*^2^ and *R_u_*^2^ showed no group differences (***Figure 3—figure Supplement 3***). By contrast, high-level visual features showed reliable group effects in face- and social-perception related regions including FFC, STS, STG, and inferior frontal cortex. The effects were largely consistent across both *R*^2^ and *R_u_*^2^ (***Figure 3—figure Supplement 4***), indicating robust differences in high-level visual information that are not explained by variance shared with low-level features.

### Exploratory: Whole-brain parcel-wise visual encoding differences and ASD±ADHD stratification

Whole-brain low- and high-level stacked encoding model analyses identified additional parcels beyond the a priori ROIs with significant visual encoding differences. At the 40% FD threshold, low-level visual *R_u_*^2^ was greater in the autism group in left 8BL (lateral BA8, posterior superior frontal gyrus; DLPFC; *q <* 0.005) and right SFL (*q <* 0.05); across thresholds, left 8BL also showed higher high-level preference in the non-autistic group (***Table 2***; cortical maps in ***Figure 5***). Other parcels with significant visual *R*^2^/*R_u_*^2^ differences at one or more thresholds included right TE1p, right PF, left IFSa, left STSvp, and striatal regions CAU-DA-lh and CAU-body-lh (***Table 2***).

**Figure 5.**
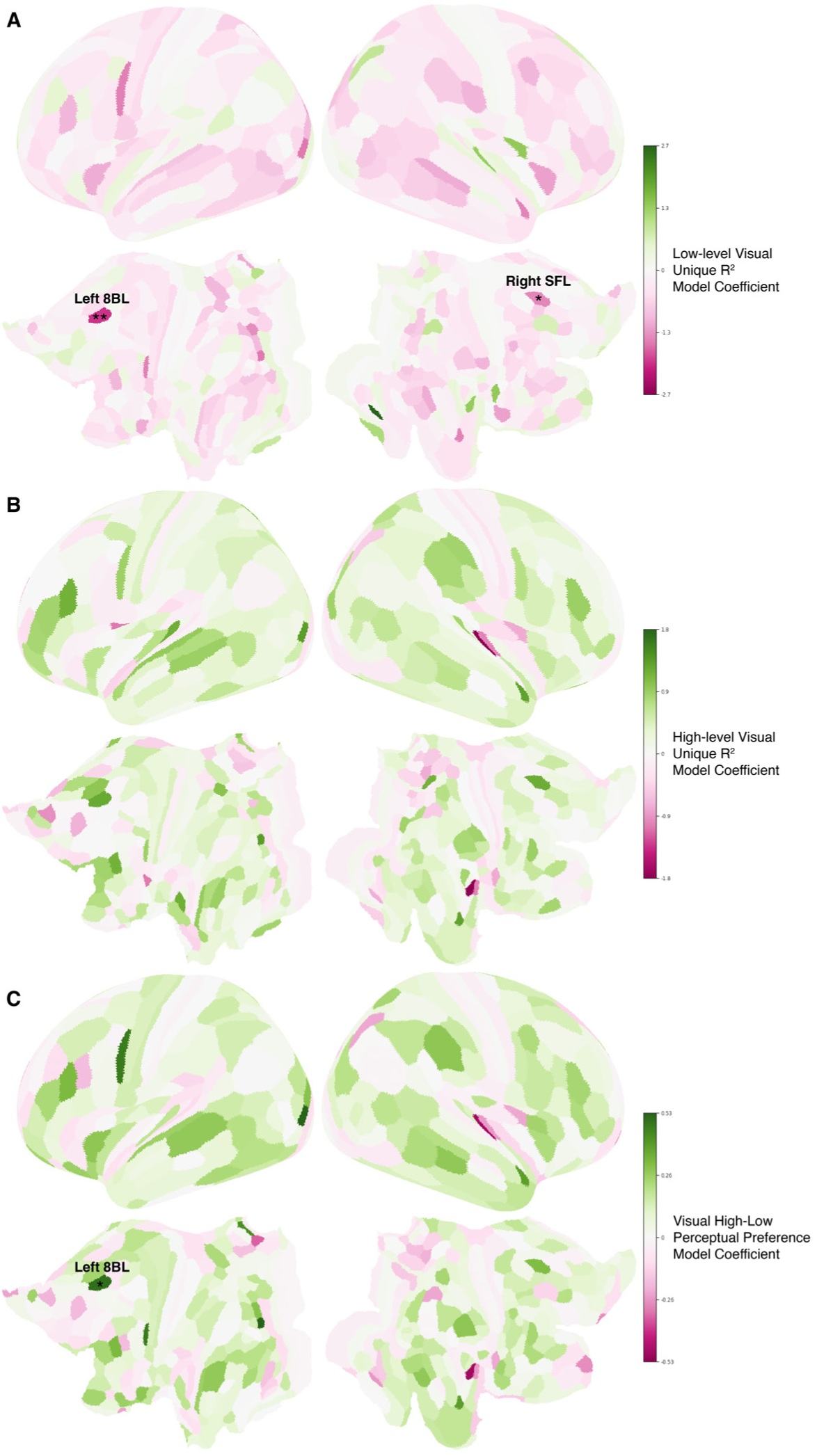
Cortical surface maps showing visual encoding model-derived fixed-effect coefficients for diagnostic group comparisons (ASD vs. nonASD) at the 40% FD threshold. Colors represent coefficient magnitude and direction (pink: negative effect, ASD > nonASD; green: positive effect, ASD < nonASD). Asterisks denote significance after FDR correction (*: FDR *q <* 0.05, **: FDR *q <* 0.005).

**Table 2.**
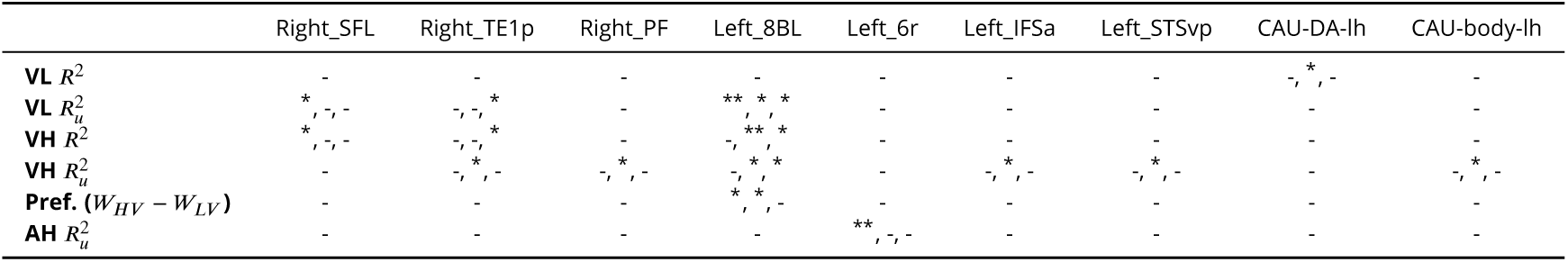
Table of post-hoc whole-brain significance results. Only significant regions and features (after FDR correction) are included in the table. In each column, results from FD thresholds of 40%, 60%, and 80% are separated by commas. VL=low-level visual, VH=high-level visual, AH=high-level audio, *W_V L_*=low-level visual model weight, *W_V H_* =high-level visual model weight, *R*^2^=explained variance, *R_u_*^2^=unique explained variance, -=q>0.05, *=q<0.05, **=q<0.005

Given the high ADHD comorbidity in the sample (and in autism generally), the autism group was further subdivided into ASD+ADHD and ASD−ADHD subgroups. Across perceptual ROIs, significant differences in visual encoding metrics were observed between nonASD and ASD−ADHD participants, but not between nonASD and ASD+ADHD (note the nonASD control group has no participants with a diagnosis of ADHD), and there were no differences between ASD subgroups (***Table 3***). The ASD−ADHD vs. nonASD results largely mirrored the broader autistic group at the ROI level as well as at the whole-brain level (e.g. a significant difference in left 8BL). A robust auditory model finding also emerged at the whole-brain level in the left Middle Insular area (MI) for low-level audio *R_u_*^2^ at 40% FD in ASD+ADHD vs. nonASD.

**Table 3.**
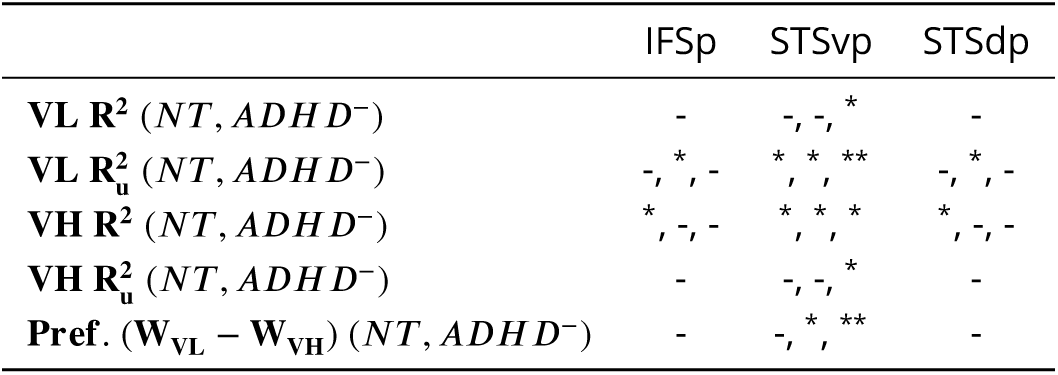
Perceptual regions of interest ADHD subgroup significance summary. All effects listed here are between the nonASD and ADHD+ASD subgroups, as there were no significant differences between nonASD and ADHD-ASD and ADHD+ASD and ADHD-ASD. Only the three regions with significant effects after FDR correction are included. In each column, the results from FD thresholds of 40%, 60% and 80% are separated by columns. VL=low-level visual, VH=high-level visual

### Exploratory: pSTS left-hemisphere predominance with ASD±ADHD corroboration

Given converging effects in the pSTS, hemispheric laterality in that region was examined in a posthoc analysis. For low-level visual features, group differences in *R_u_*^2^ were evident in right STSvp but not left (***Figure 6*** A). In contrast, for high-level visual *R_u_*^2^ and *W_H_* − *W_L_*, group differences were stronger and more consistent in the left hemisphere (STSvp and STSdp; ***Figure 6*** B,C). Right STSvp trended in the same direction for both *R_u_*^2^ and *W_H_* − *W_L_* but did not survive FDR at the 40% FD threshold (uncorrected *p < .*05). Taken together, this pattern suggests a hemispheric division of effects. ADHD-stratified analyses corroborated this laterality. Comparing ASD−ADHD with nonASD reproduced the pSTS effects with the same left-hemisphere predominance across metrics (***Figure 6*** D-F)).

**Figure 6.**
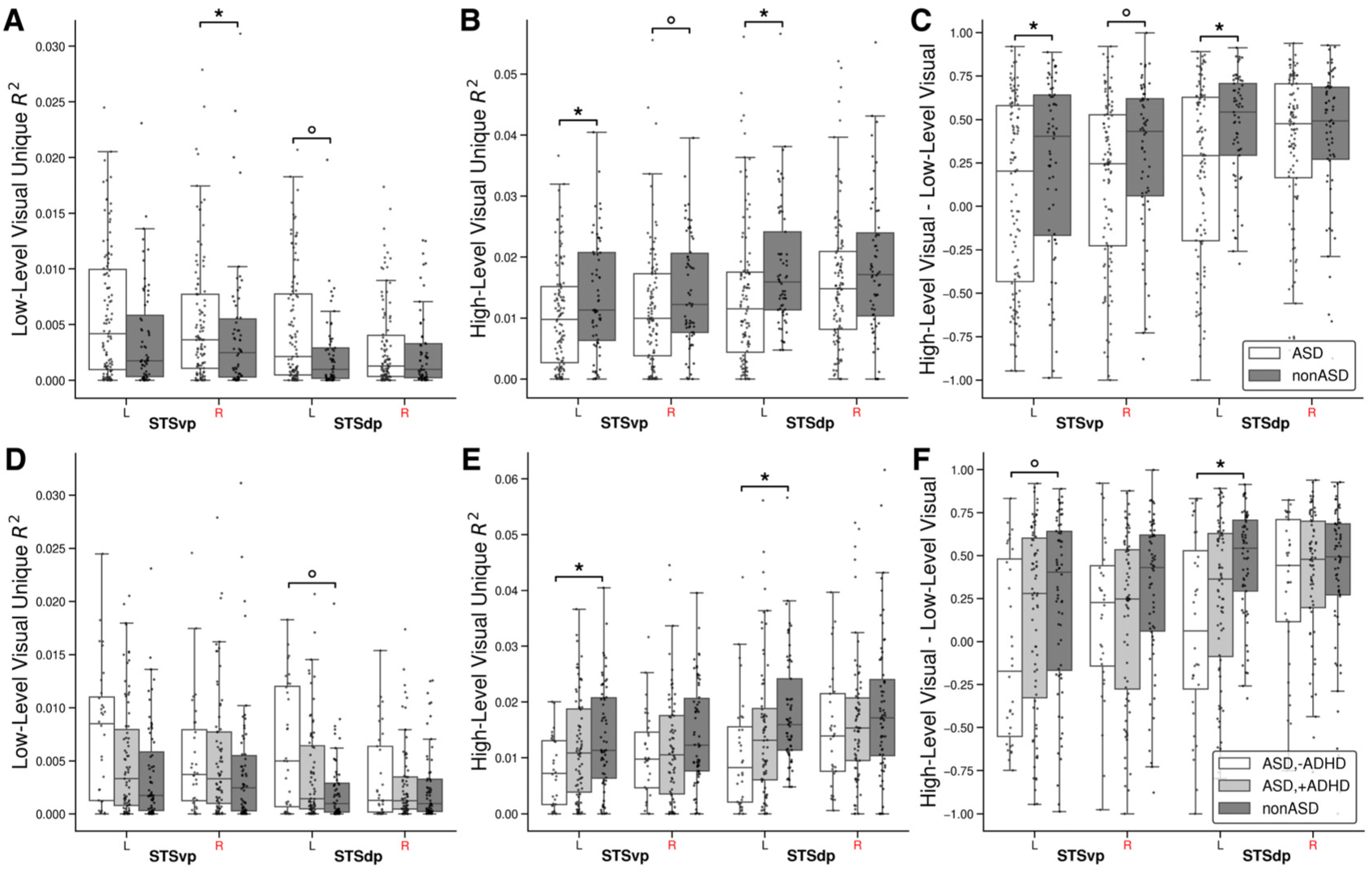
Visual encoding model metrics across left and right pSTS subregions. (A) Low-level visual unique explained variance (*R_u_*^2^) for ASD (white) and nonASD (gray) groups in left and right STSvp and STSdp regions. (B) High-level visual *R_u_*^2^ across the same groups and regions. (C) High- vs. low-level perceptual preference (derived from the stacked visual encoding models), where positive values indicate a high-level preference, negative values indicate a low-level preference, and values near zero indicate no preference. (D) Low-level visual *R_u_*^2^ for ASD-ADHD (white), ASD+ADHD (light gray), and nonASD (gray) groups across left and right STSvp and STSdp regions. (E) High-level visual *R_u_*^2^ across groups in the same regions. (F) Stacked encoding model visual perceptual preference metrics (high- vs. low-level); positive values indicate a preference for high-level features, negative values indicate a low-level preference, and values near zero indicate no strong perceptual preference. All results are from the 40% FD threshold. Boxes annotated with an asterisk (*) indicate significant group differences (FDR-corrected *p <* 0.05); circles indicate nominal significance prior to FDR correction (uncorrected *p <* 0.05). Boxes represent quartiles; whiskers indicate data range excluding outliers. Individual dots represent mean metrics per subject, averaged across statistically significant grayordinates within each left or right ROI. Pairwise significance tests were conducted between all groups; annotations indicate significant differences, all of which occurred between the ASD-ADHD and nonASD subgroups.

### H2: Audio–visual feature representation and modality preference is conserved across groups and not related to autism severity

There were no significant ASD vs. nonASD differences in visual vs. audio preference (*W_V_* − *W_A_*, ***Figure 7***) or audio and visual *R*^2^ (***Figure 7—figure Supplement 1***) or *R*^2^ (***Figure 7—figure Supplement 2***) within their corresponding cortical regions, with the singular exception of TPOJ2 at the 60% FD threshold (greater visual *R*^2^ in the nonASD relative to ASD group, ***Figure 7—figure Supplement 3***). Overall we observed minimal evidence of reduced visual dominance (or increased auditory dominance) related to audio and visual feature representation in autism during naturalistic viewing. Beyond diagnostic group differences, audio and visual model *R*^2^, *R_u_*^2^ and *W_V_* − *W_A_* were also not significantly related to SRS or SSS (***Figure 7—figure Supplement 3*** B,C).

**Figure 7.**
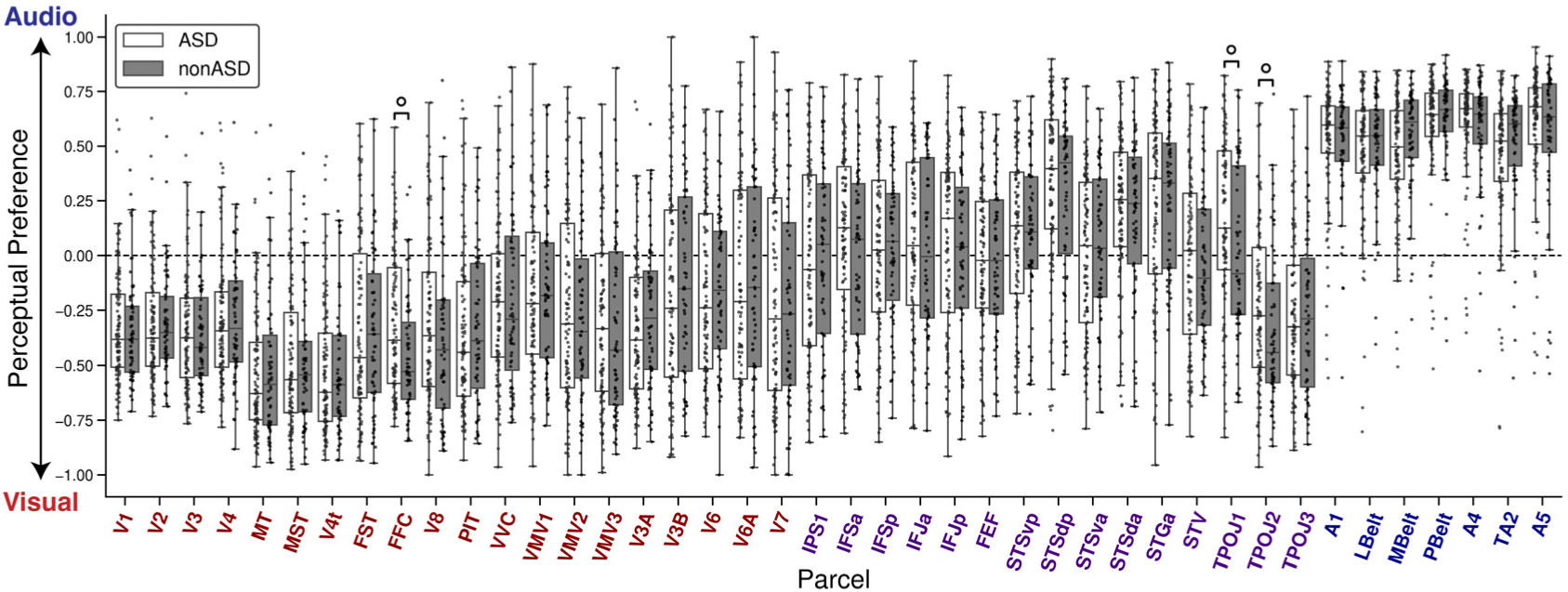
Audio vs. visual modality preference. Higher values indicate an audio preference and lower values indicate a visual preference. These results correspond with the 40% FD threshold. Boxes annotated with circles indicate a difference between groups that did not survive FDR correction (uncorrected p<0.05 but FDR q>0.05). Boxes show the quartiles of the dataset and whiskers show the distribution with the exception of outliers. Each dot is the mean value from a single subject from all statistically significant grayordinates within each ROI.

### Exploratory: Age exerts prominent effects on modality preference

Focusing on age without regard to diagnostic group, robust relationships emerged across perceptual ROIs (***Figure 8***). Visual *R*^2^ and *R_u_*^2^ increased with age in V3, MT, MST, V3A, IFSa, TPOJ3, and A4; auditory *R*^2^ and *R_u_*^2^ increased with age in VMV2, V3A, IFSa, IFSp, IFJa, STSdp, STGa, TPOJ1, PBelt, A4, TA2, and A5. Modality preference (*W_V_* −*W_A_*) shifted with age in V3A, IFSa, IFSp, A4, and A5, such that the visual region V3A had an increase in visual preference and the other auditory and multisensory regions had an increased auditory preference. Notably, these age effects were prominent at the 60% and 80% FD thresholds but not at the 40% threshold, plausibly because this stricter criteria disproportionately excluded more younger participants due to their increased motion.

**Figure 8.**
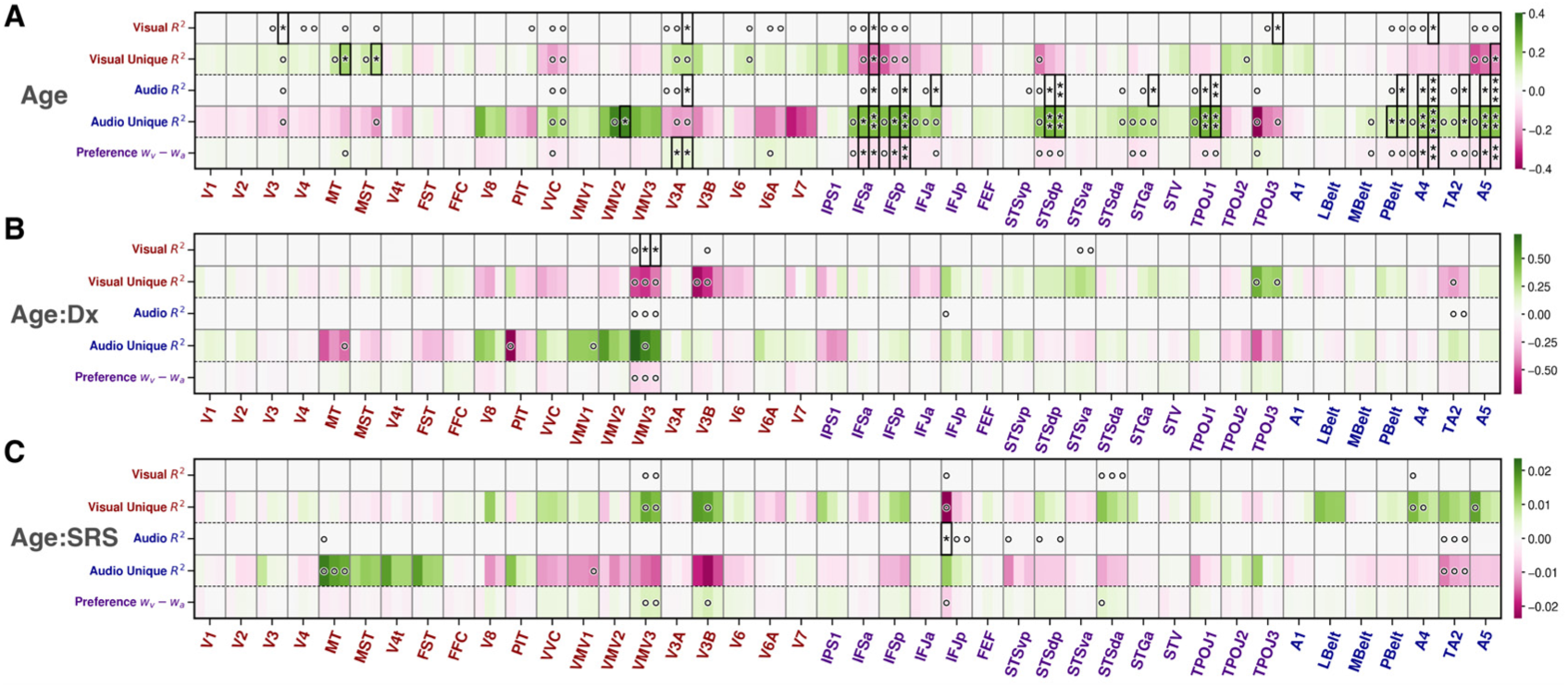
Heatmaps of fixed-effect coefficients from exploratory parcel-wise linear mixed models examining (A) main effects of age, (B) age-by-diagnosis (ASD vs. nonASD), and (C) age-by-SRS interactions. Within each horizontal panel, rows represent encoding model-derived metrics (visual *R*^2^ and *R_u_*^2^, audio *R*^2^ and *R_u_*^2^, and their perceptual preference index *W_V_* − *W_A_*), while columns represent Glasser MMP parcels ordered roughly from early visual through higher-order association to auditory areas. Each parcel is further subdivided by FD thresholds (40%, 60%, 80%, left to right). Color indicates magnitude and direction of coefficients (green: positive effects; pink: negative effects). In panel A, green indicates encoding metrics increase with age, pink indicates metrics decrease with age. In panel B (Age × Diagnosis), green indicates the age-related slope is more positive in nonASD (greater increases with age), whereas pink indicates a more positive age-related slope in ASD. Asterisks mark FDR-corrected significance (∗= *q <* 0.05,∗∗= *q <* 0.005,∗∗∗= *q <* 0.0005); open circles mark uncorrected significance (*p <* 0.05). Visual metrics are labeled in red; auditory metrics in blue.

Whole-brain analyses (where all cortical and subcortical parcels were considered separately across the left and right hemispheres) confirmed many ROI effects and further revealed some hemispheric specificities (***Table 4***): for example, auditory *R*^2^ related to age in A5 bilaterally, whereas V3A/MT effects were right-lateralized and A4/STSdp/TPOJ1 were left-lateralized. Beyond the predefined perceptual ROIs, additional parcels with significant age relationships included bilateral LO3, right TGv, left 9a, and left RI, adjacent to primary auditory cortex (***Glasser et al., 2016***). Several language-network parcels (pSTS/55b/PSL/SFL/area 45) also showed age effects for auditory *R*^2^ and *R*^2^, predominantly left-lateralized (***Glasser et al., 2016***). In all cases, the directionality matched modality (auditory metrics increased with age in auditory regions; visual metrics increased with age in visual regions), and modality preference became more congruent with regional specialization (See ***Figure 8—figure Supplement 1*** and ***Figure 8—figure Supplement 2*** for examples from perceptual and whole-brain regions respectively).

**Table 4.**
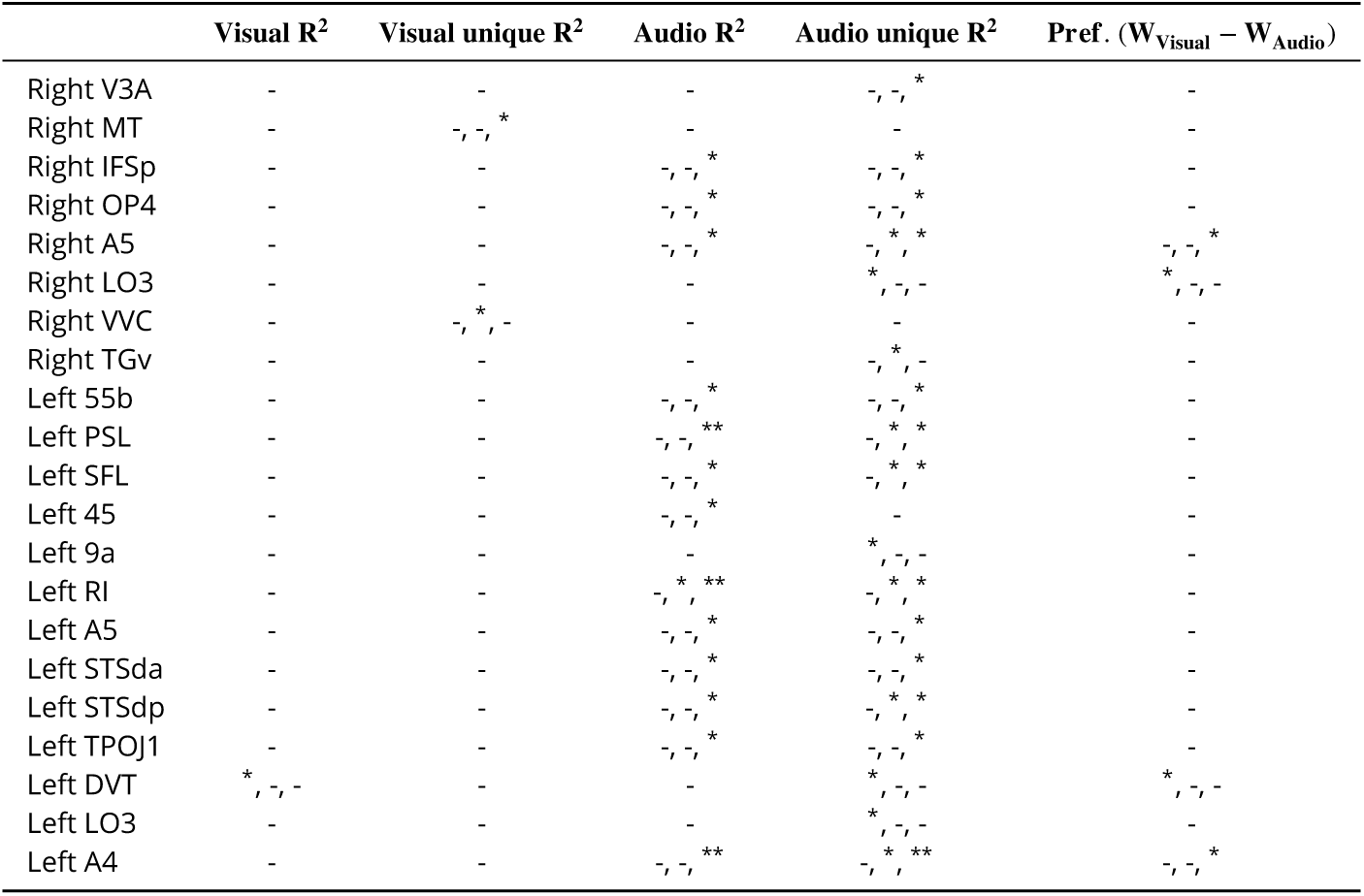
Significance of Age effects on visual, audio, and audio–visual encoding model metrics across all left and right hemisphere, cortical and subcortical parcels at the whole-brain level.

Post-hoc whole-brain ADHD subgroup comparisons revealed one region with consistent auditory *R*^2^ differences that survived correction across all 410 parcels: left PoI2 (insula), with ASD−ADHD vs. noNASD differences across all three FD thresholds (FDR *q* = 0.03, 0.003, 0.008 for 40%, 60%, and 80%, respectively). No other pairwise subgroup contrasts (ASD−ADHD vs. ASD+ADHD; ASD+ADHD vs. nonASD) yielded auditory effects that survived FDR correction.

### Exploratory: Age-by-diagnosis and Age-by-SRS interactions are sparse and localized

At the ROI level, only VMV3 showed a significant Age×Diagnosis interaction for visual *R*^2^ where in the autistic group, visual *R*^2^ was relatively stable across age, whereas in the non-autistic group it decreased with age (***Figure 8—figure Supplement 3***). Whole-brain analyses identified two additional Age×Diagnosis interactions for visual *R_u_*^2^: right area 52 and left p32pr, where visual *R_u_*^2^ increased with age in autism but decreased with age in non-autistic participants. For Age×SRS, only audio *R*^2^ in IFJp showed a significant interaction at the ROI level (autism: slight increase with age; non-autism: decrease), and at the whole-brain level, interactions appeared for visual *R*^2^ and *R_u_*^2^ in right 7PL (the autistic group measure slightly decreasing with age while the non-autistic group measure increases with age).

### Results Summary

Overall, this study finds (i) no evidence for increased low-level encoding specific to primary sensory cortices in autism; (ii) reduced high-level visual feature representations in social and face brain regions (STSvp, STGa, FFC) in autism; (iii) a relative preference towards low-level visual features in the pSTS in autism, strongest in the left hemisphere and linked to SRS score; and (iv) conserved audio–visual modality dominance and feature encoding in autism. Further, results show that feature encoding becomes increasingly more modality-congruent with age, with lateralized parcel-specific trajectories, though interactions of age with diagnosis or SRS score are sparse. Post-hoc whole-brain and ADHD-stratified analyses further identify additional brain regions with significant encoding differences, including left 8BL and left PoI2.

## Discussion

We asked where and how sensory features are differentially represented in the brain during naturalistic viewing in autism. First, we found no evidence for stronger low-level encoding in primary sensory cortices in autism; instead, differences localized to higher-order association regions, most notably the pSTS and adjacent frontal parcels. Second, in the autistic group, pSTS showed a preference for lower-level visual information over high-level visual encoding, with perceptual preference significantly related to SRS scores. Third, audio–visual modality balance was broadly conserved in autism, with significant effects predominantly observed developmentally in relation to age rather than phenotypically across diagnostic groups.

### Implications for sensory theories

The observed pattern of downstream, visual-specific alterations, with reduced high-level representations in social and integration hubs, fits the theory of weak central coherence (WCC) more readily than enhanced perceptual functioning (EPF), which predicts increased encoding in early sensory cortex. At a minimum, high-level global integration differences appear to outweigh any enhancements of early visual coding in this dataset and experimental context. A hybrid account remains plausible, where both theories are true or where subtle early differences cascade to association regions, for example, but our results do not directly support this.

### pSTS, lateral pathway, and social perception

The emergence of the pSTS as a locus of autistic group differences in our results is mechanistically informative. The pSTS is known to support perceptual functions including audiovisual speech (***Okada et al., 2013***), biological motion (***Sokolov et al., 2012***), and higher-order social perception (***Lahnakoski et al., 2012***), with changes observed across development (***Xiao et al., 2016***). Here, autistic participants showed decreased high-level visual encoding and a relative preference toward low-level visual features, with a left-hemisphere predominance for the high-level reduction. This lateralized pattern sits within the context of the proposed third, lateral visual pathway thought to be specialized for social perception, and raises the possibility that altered visual components of socially relevant signals might underlie reported downstream effects, such as reduced audiovisual speech perception, observed behaviorally in autism. One key detail is that the SRS association was specific to the perceptual preference measure (*W_H_* −*W_L_*) rather than raw performance, pointing to an altered weighting of visual evidence rather than a uniform gain/loss. Another key observation is that high-level visual *R*^2^ itself was reduced in social brain regions (FFC/STS/STG), consistent with a selective reduction in category-level visual information and not a broader effect that might be attributable to something like increased noise.

### Audio–visual dominance: largely conserved

Contrary to predictions related to behavioral “reverse Colavita” reports, we did not observe group differences in audio–visual modality preference in sensory cortices. Naturalistic, passive viewing and ceiling/floor constraints in unimodal cortices may attenuate sensitivity to cross-modal shifts behaviorally measured under different contexts; moreover, the sensory dominance transitions reported in psychophysics may precede our sampled age range or require task demands that are absent here. An interpretation of these findings is that shifts in sensory modality dominance, at least at the levels measurable by fMRI-based encoding models, are not a hallmark of autistic perception during naturalistic movie viewing.

### Developmental effects dominate audio and visual encoding

Age explained more variance than the diagnostic group when considering the stacked audio-visual encoding models. Specifically, encoding performance increased with age within corresponding modalities (visual model in visual cortex; audio model in auditory cortex, multisensory, and language areas), and modality preference also became more congruent with regional specialization. Effects were often lateralized (e.g., right dorsal-stream visual parcels; left language network), consistent with known specializations and a general network-specific reweighting. These results demonstrate the ability of encoding models to pick up broad developmental effects while ruling out these same factors in relation to diagnosis and phenotype. Notably, most age-related effects appeared only at looser FD thresholds, and stricter FD thresholds disproportionately excluded younger participants. This highlights the importance of performing analyses with an awareness and transparency around factors such as noise and data quality, as this can be a critical factor that affects downstream results and inferences.

### ADHD comorbidity

Post-hoc stratification results suggest that visual encoding differences are driven by autistic participants without ADHD; the ASD+ADHD subgroup showed little divergence from the non-autistic group, with a lone insular auditory effect under the strictest motion threshold. This pattern motivates explicit modeling of ADHD comorbidity in future work and raises the possibility of potentially opposing sensory feature encoding in ASD vs. ADHD. This finding is particularly interesting, given that functional connectivity has been shown to predict SRS across both ADHD and ASD (***Lake et al., 2019***).

### Sex-based considerations

Autism shows sex-linked differences in prevalence, diagnostic presentation, and neurobiology, with growing evidence for partly distinct neural phenotypes in females and males (***Napolitano et al., 2022***). Although sex was included as a covariate, the present study was neither designed nor powered to adjudicate sex-specific encoding effects. Consequently, the pSTS findings and high– vs. low-level visual reweighting could be disproportionately driven by males. Future work should pre-register sex-stratified analyses (and Sex × Diagnosis and Sex × Age interactions), recruit adequately powered female cohorts, and incorporate pubertal status as a moderator. Such designs will clarify whether the observed reductions in high-level visual encoding within social–perceptual networks reflect a male-typical autistic phenotype, a shared mechanism across sexes, or qualitatively different pathways in autistic females.

### Limitations and Future Work

- Stimulus duration. The combined movie runs were comparatively brief, which limited encoding performance, constrained the feasible feature space (due to overfitting when increasing the number of features), and limited the ability to perform noise-ceiling and within-subject reliability estimation. We therefore used compact, theoretically robust features with crossvalidation and permutation testing. Future work using longer acquisitions (and/or repetitions) would permit richer models to probe for finer-grained effects.
- MSI-related inferences are limited by modality and design. With only mixed audiovisual runs, no unimodal (A-only/V-only) acquisitions, and no behavioral MSI measures (e.g., temporal binding window or Colavita task), direct tests of multisensory integration were not possible. Moreover, many audiovisual interactions unfold at millisecond scales beyond the resolution of the BOLD signal. We therefore used modality-resolved encoding and variance partitioning to separate audio and visual contributions within fMRI’s constraints. Future work should incorporate unimodal runs and MSI tasks, and pair fMRI with EEG/MEG to capture fast crossmodal dynamics.
- Sensory phenotyping. No dedicated sensory-processing scales (e.g., SSP/SPM) were available to map our results directly to sensory phenotypes. Using the scales available in the dataset with sensory related questions (SRS-2 and SCQ) we aggregated and used a Sensory Subset Score (SSS), though it was not psychometrically validated. Future work should pair encoding model results with dedicated sensory-processing scales or validate the SSS.
- Sample composition and generalizability. The dataset included only verbally fluent participants and the autism cohort had a male-skewed sex distribution and high ADHD comorbidity, limiting generalizability, particularly excluding minimally and non-verbal autistic individuals and autistic females. We included age, sex, socioeconomic status, and site as confounds in all analyses, and in exploratory analyses, stratified by ADHD diagnosis, but future work should balance sex, ADHD, and verbal ability and model comorbidity a priori.
- Preprocessing/parcellation choices and lack of functional localizers. We used MMP parcels without full MSMAll alignment and without subject-specific functional localizers (e.g., a face localizer for FFA) as they were not available. This may have introduced some degree of registration imprecision and ROI mis-localization. We did perform MSMSulc cortical surface registration, allowing for improved surface alignment. Future work could apply MSMAll alignment, functional localizers, or subject-specific parcellations to improve anatomical specificity.
- Eye gaze and attention. Autistic differences in fixation patterns during naturalistic viewing are well documented and can alter effective visual input (***Klin et al., 2002***). As eye-tracking data was not directly collected during fMRI acquisiton and regressors were not gaze-weighted, we cannot exclude eye-movement confounding or that our observed differences may be due to autistic participants watching the movie differently. This warrants future work incorporating eye-tracking and gaze-weighted features.
- Network-level dynamics and directionality. Our analyses were regional and did not assess how feature representations are coordinated across networks or the direction of information flow. Within these constraints we limited inference to parcel-wise metrics. Future work should relate encoding to static and time-varying functional connectivity and apply effectiveconnectivity approaches to test directionality, with pSTS and adjacent social–integration hubs as a priori targets.

## Conclusion

In conclusion, through preregistered stacked encoding model analyses applied to naturalistic movieviewing fMRI data from the Healthy Brain Network, we find little support for pervasive early sensory enhancement accounts of autism at the level of feature-resolved cortical representations. Instead, autism-related differences are concentrated in higher-order integration and social–perceptual regions, most notably the pSTS, where high-level visual representations are reduced and feature weighting shifts toward lower-level visual information in a manner that tracks social symptom severity. At the same time, audio–visual modality preference and unimodal dominance appear largely conserved, arguing against large global shifts in sensory dominance during passive naturalistic viewing. Developmental effects were prominent, lateralized, and modality-congruent, underscoring the need to interpret group differences against ongoing maturation and motion-related sampling constraints in pediatric datasets. Together, these results support mechanistic interpretations related to altered visual feature weighting within social/multisensory networks and demonstrate that naturalistic stimuli coupled with encoding-models can provide a scalable, theory-relevant framework for characterizing how neurodevelopmental conditions may reshape sensory representations in the brain.

## Methods

### Existing Data

This study is based on two pre-registrations (accessible at osf.io/h92gr and osf.io/47kj6; summarized in ***Table 1***) relying on data from releases 1-10 from the openly accessible Healthy Brain Network (HBN) dataset from the Child Mind Institute (***Alexander et al., 2017***). The HBN dataset offers a large cross-sectional population that is highly phenotypically characterized. The combination of a young population, naturalistic imaging acquisitions, and scales like the social responsiveness score make it an ideal dataset for this analysis.

#### Explanation of Existing Data

As additional data is released, a growing body of prior research has analyzed the HBN dataset (***Kember et al., 2024***; ***Mihailov et al., 2020***; ***Tong et al., 2024***; ***Cohen et al., 2022***; ***Camacho et al., 2023***; ***Di et al., 2023***). To our knowledge, at the time of this writing, there is no existing research using naturalistic fMRI to predict autism diagnostic status and SRS score from this dataset. Some metadata relevant to the research plan including summary statistics, phenotypic data and quality control metrics from the fMRI data were accessed and analyzed prior to the pre-registration of this work. As part of a pilot study to select features and set thresholds and methodological parameters for the pre-registrations, functional data from 54 participants who are neither in the autism nor control groups were accessed. These participants were withheld from all subsequent analyses.

### Behavioral Measures

#### Autism diagnosis

Autism diagnosis relied on clinically synthesized consensus DSM-5 diagnoses. These diagnoses were generated by the HBN clinical team based on interviews including the Schedule for Affective Disorders and Schizophrenia—Children’s version (KSADS) and all materials and questionnaires collected during study participation in addition to behavioral observations, family history, previous diagnoses, and history of therapeutic intervention. A limited subset of participants with autism-related traits were administered the Autism Diagnostic Observation Schedule (ADOS). All phenotypic data were accessed in accordance with a data usage agreement from the Child Mind Institute.

#### Social Responsiveness Scale (SRS-2)

The SRS is a quantitative measure of autistic traits that is correlated with other commonly used scales including the ADOS and ADI-R (***Bölte et al., 2008***; ***Constantino et al., 2003***; ***Constantino, 2007***).

#### Sensory subset score

While we aim to link neural signatures to targeted behaviorally measured sensory differences in autism, assessments that specifically target such differences, like the Short Sensory Profile (SSP), were not included in the HBN study. To overcome this limitation, a sensory subset score (SSS) was generated from 3 items on the SRS-2 and one item from the Social Communication Questionnaire (SCQ) (***Rutter et al., 2003***), both of which all participants took part in. While the SRS-2 is a well-validated measure, the SSS is not psychometrically validated, but may be more specific to sensory-related differences. SSS is composed of the sum of items 20, 42, and 58 of the SRS-2 and item 14 of the SCQ relating to unusual sensory interests, sensory oversensitivities, and concentrating on parts instead of the whole. Specifically, these are: *“I have sensory differences that others find unusual (for example, smelling or looking at things in a different way).”, “I am overly sensitive to certain sounds, textures, or smells.”, “I concentrate too much on parts of things rather than seeing the whole picture.”,* and *“Has he/she ever seemed to be unusually interested in the sight, feel, sound, taste, or smell of things or people?”*. SSS is a sum of these scores where 0=No/Not True, 1=Sometimes True, 2=Often True, and 3=Almost Always True/Yes with the minimum SSS score being 0 and the max being 12.

### Group-level Confounds

All primary analyses will be conducted at the individual level before demographic confounds are accounted for. Demographics including age, sex, and socioeconomic status along with site-level variability will be included as confounds in all group-level statistical analyses. Family socioeconomic status will be reported as the total score from the Barratt Simplified Measure of Social Status (BSMSS).

### fMRI data

The analysis focuses on two naturalistic movie-viewing fMRI acquisitions (*Despicable Me*, 10 min; *The Present*, 4 min). All fMRI data (TR=0.8s) were acquired from 3 scanner sites in the Greater New York area.

### Quality Control Criteria

Motion-based exclusion criteria in neuroimaging studies of autistic children is an important consideration (***Elyounssi et al., 2025***) and quality control criteria have been shown to alter the distribution of included subjects. Specifically, subjects that are younger and have higher ADOS and SRS scores, along with more inattentive, hyperactive/impulsive, and motor control features are more likely to be excluded due to motion (***Nebel et al., 2022***). To account for this issue, all results are reported across a sweep of 3 framewise displacement (FD) based functional motion thresholds. Specifically, functional data is excluded if the number of time points in a run with a mean FD > 0.2mm is greater than 40%, 60%, or 80%. By starting with a more stringent threshold and sweeping this across more lenient parameters, we aim to strike a balance between the best quality and more fully characterizing as much of the dataset as possible. In this sample specifically, head motion (fraction of volumes with FD > 0.2 mm) decreased with age (Spearman *ρ*= -0.40), but showed weak linear associations with SRS Total T and SSS (*ρ* = -0.01 and 0.06, respectively), which remained small after controlling for age (partial *ρ* = 0.03 and 0.08). As neither SRS nor SSS were correlated with fractional FD, this pattern suggests that age is a primary driver of motion in this cohort and not symptom severity (see ***Figure 9***).

**Figure 9.**
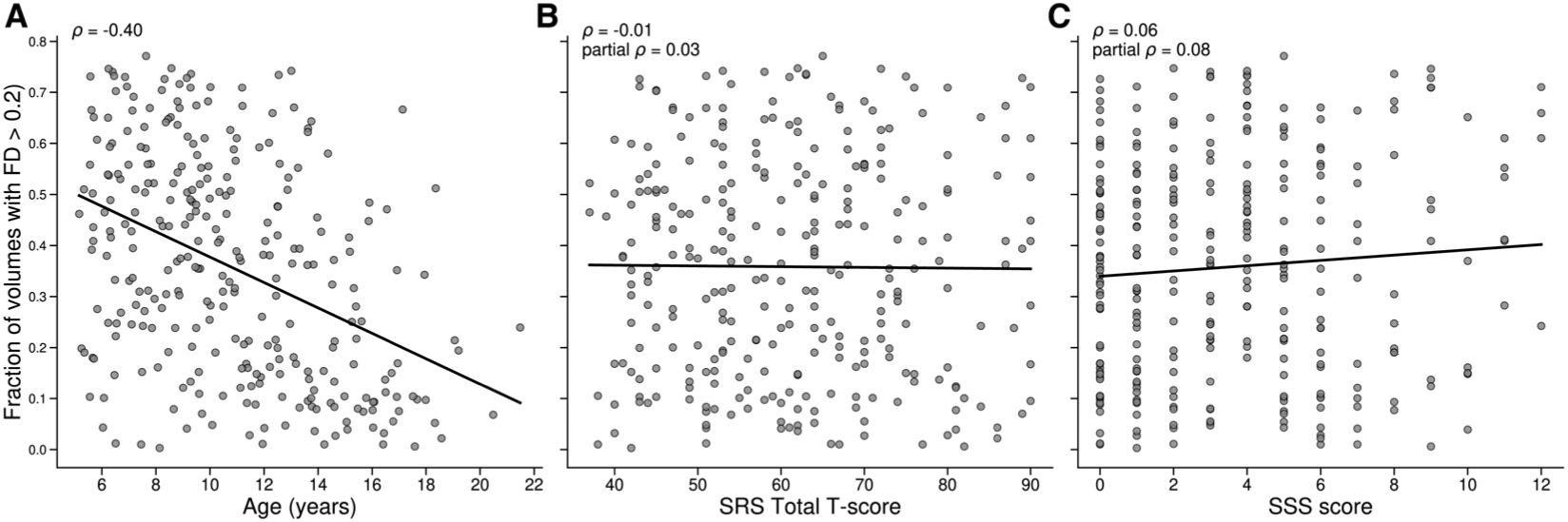
Head motion vs. age and behavioral measures. Scatterplots show the fraction of volumes with framewise displacement *>* 0.2 mm in relation to (A) age, (B) SRS Total T-score, and (C) Sensory Subset Score (SSS). Solid black lines denote linear fits. Reported coefficients are Spearman correlations (*ρ*); “partial *ρ*” controls for age. (A) *ρ* = −0.40; (B) *ρ* = −0.01, partial *ρ* = 0.03; (C) *ρ* = 0.06, partial *ρ* = 0.08. Motion decreases with age and shows only weak associations with SRS/SSS.

Quality control is not limited to functional data and is also applied to T1 data. T1 quality is assessed in a semi-automated procedure via a Bayesian DNN brain parcellation method which provides an uncertainty metric highly correlated with T1 structural image quality (***McClure et al., 2019***). Images with mean uncertainty *>*= 0.4 are flagged for manual inspection of their structural images and subjects with excessive artifacts and irregularities are excluded from all analyses. In addition to these semi-automated approaches, a manual QC evaluation is performed by examining the reconstructed surfaces both alone and overlaid on subject T1 images across all subjects. This entire process is performed “blind”, i.e. all visual inspections are performed without knowledge of which cohort a subject belongs to or any other demographic information.

### Diagnostic Inclusion Criteria

A subset of the total subjects (n=4767 before quality control and before excluding pilot subjects) are included in analyses based on their diagnostic status as determined by the clinician consensus field from the phenotypic and demographic data. Subjects in the autism group are selected if they have received the DSM-5 label of ‘Autism Spectrum Disorder’ (n=659 before quality control) and non-autistic participants, referred to here as nonASD, are selected if they have the label ‘No Diagnosis Given’ (n=373 before quality control). Individuals with the labels ‘No Diagnosis Given: Incomplete Eval’ and ‘No Diagnosis Given: No Reason Given’ are excluded from all analyses. Any individuals with co-occurring conditions including ADHD are not excluded from the autism group, despite the significant overlap (n=176 autism only, n=483 both autism and ADHD) in order to fully sample the phenotypic space of autism present in the dataset. Additional secondary analyses are performed to compare the autism only group to controls.

### Demographics

After quality control and filtering (see ***Figure 10*** for a flow chart) at the most stringent FD threshold, 63 non-autistic participants (mean ± SEM age = 10.96 ± 0.46 years; WISC–FSIQ = 110.8 ± 2.05; Barratt Total = 54.4 ± 1.39; SRS Total T = 49.3 ± 0.97; SSS = 1.25 ± 0.21) and 108 autistic participants (mean ± SEM age = 12.76 ± 0.37 years; WISC–FSIQ = 101.4 ± 2.08; Barratt Total = 49.3 ± 1.38; SRS Total T = 68.1 ± 1.10; SSS = 4.71 ± 0.28) were retained. Sex distribution differed significantly between groups (*χ* ^2^(1) = 14.23*, p* = 1.6 × 10^−4^), with a higher proportion of males in the autism cohort. Normality (Shapiro–Wilk or D’Agostino’s K²) and homogeneity of variance (Levene’s test) were assessed for each continuous measure within each diagnostic group. Where both normality and equal variance held, Student’s *t*–tests were applied; where variances differed, Welch’s *t*; and for non-normal distributions, two-sided Mann–Whitney *U* –tests were used. Sex differences were evaluated via *χ* ^2^, with Fisher’s exact test substituted if any cell count fell below five. To control the family-wise error rate over six univariate comparisons, *p*–values were adjusted via the Benjamini–Hochberg false discovery rate (*α* = 0.05). All demographic measures differed significantly between non-autistic and autistic groups after FDR correction, but a multivariate logistic regression (diagnosis ∼ Age + FSIQ + Barratt + SRS + SSS + Sex) confirmed that, when entered simultaneously, only SRS (*z* = −3.84, *p* = 1.2 × 10^−4^) and sex (*z* = 2.70, *p* = 6.8 × 10^−3^) remained significant independent predictors of diagnostic status.

**Figure 10.**
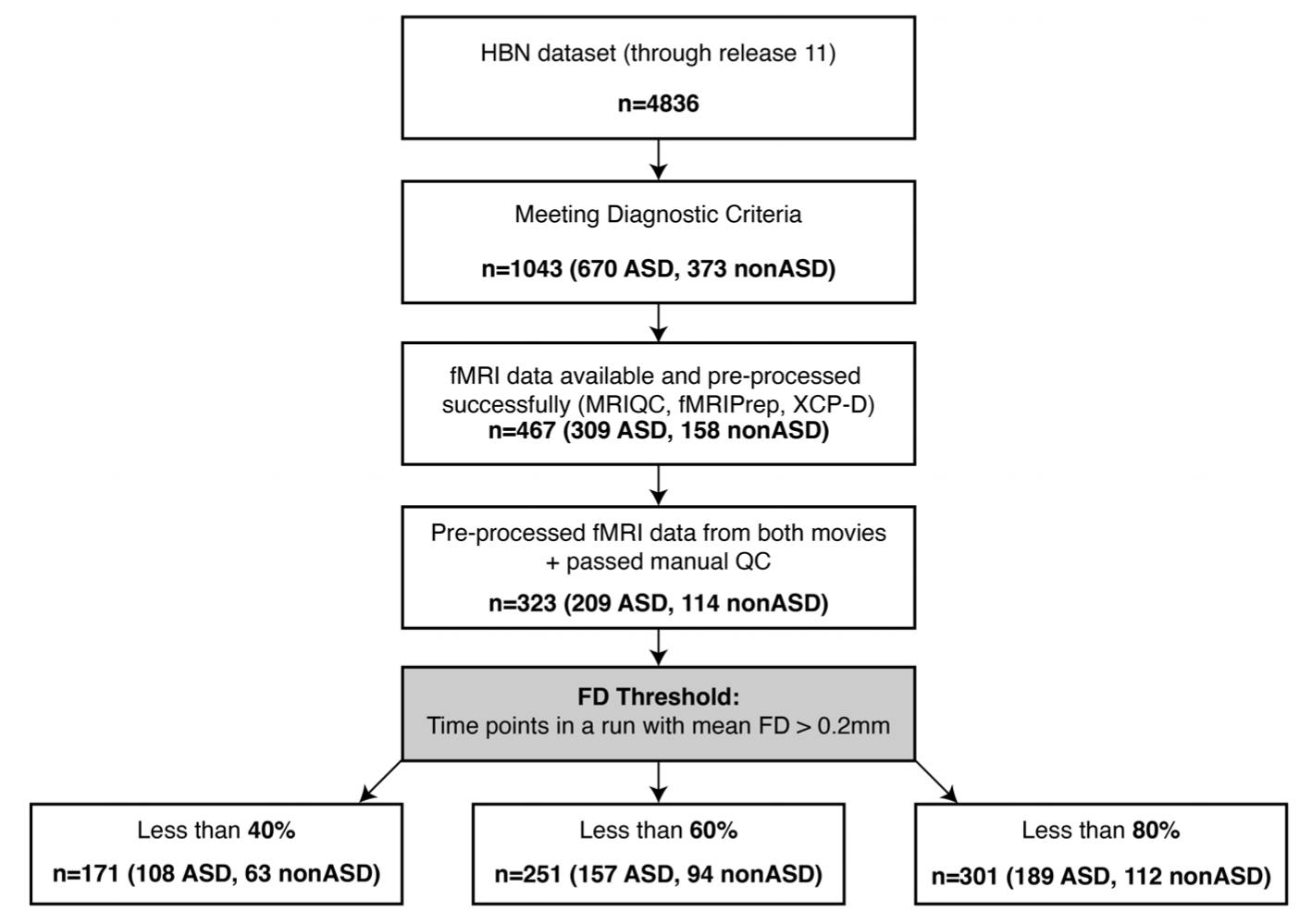
A flow chart illustrating how participants were selected for the final sample including the number that were excluded at each stage.

**Table 5.**
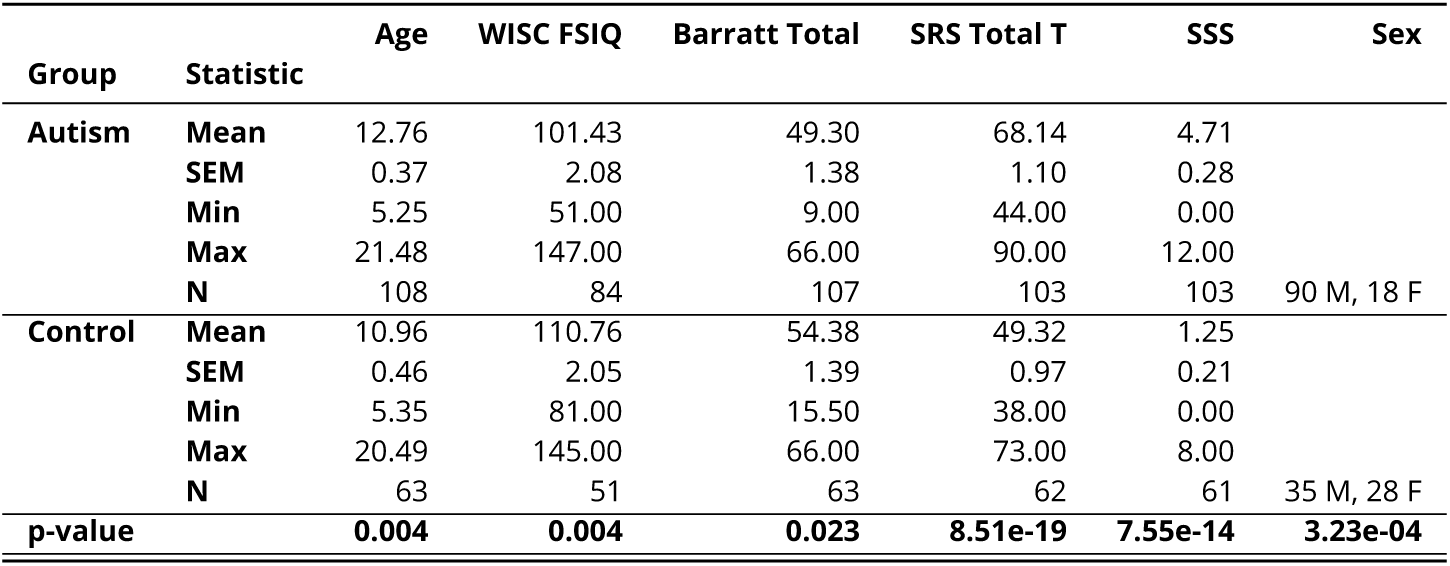
Descriptive statistics of demographic and clinical variables by diagnostic group. Continuous measures (Age, WISC–FSIQ, Barratt Total, SRS Total T, SSS) are reported along with their mean, SEM, minimum–maximum range in parentheses. “N” indicates the number of non-missing observations. Sex shows counts of males and females. P-values are FDR-corrected (Benjamini–Hochberg, *α*=0.05).

### fMRI Preprocessing

fMRI data from all included subjects was preprocessed using fMRIprep version 23.2.0 (***Esteban et al., 2019***). This version of fMRIprep includes MSMSulc cortical surface registration, allowing for improved surface alignment compared to previous versions of fMRIprep. CIFTI2 outputs (combined cortical surface and subcortical volume) were obtained in fsLR 32k space. All subsequent analyses were performed using CIFTI images to better respect the geometry of brain structures of interest and prevent issues such as smoothing across cortical folds. The first 10 timepoints from each run were discarded. Further preprocessing was performed using XCP-D version 0.10.1 (***Mehta et al., 2024***). Specific steps included: 4mm FWHM spatial smoothing, despiking, 0.1 Hz low-pass filtering, confound regression of: six motion estimates and their derivatives, five white matter, and five CSF ACompCor components, denoising and detrending.

### Parcellation Scheme

The Glasser Multi-Modal Parcellation (MMP) (***Glasser et al., 2016***) was used to select regions of interest for all analyses. Regions were selected from auditory regions and across the dorsal, ventral, and lateral visual streams.

### Feature Selection

Movie features include automatically extracted and hand-labeled low- and high-level audio and visual classes. These features were selected based on their performance on an independent set of 54 pilot subjects that were excluded from all subsequent analyses. Low-level auditory features were extracted from the audio track of the movie files and include perceptual loudness extracted via pyloudnorm (***Steinmetz, 2023***) and the first 5 principal components of the cochleagram extracted via pycochleagram (***McDermott, 2018***). AudioSet (***Gemmeke et al., 2017***) labels comprise the high-level auditory features and were extracted using YAMNet. Instead of a 521-dimensional vector of all AudioSet classes, classes were combined across the highest level of the AudioSet ontology yielding an 8-dimensional feature consisting of: ‘Human Sounds, Speech’, ‘Human Sounds, Non-Speech’, ‘Animal Sounds’, ‘Music’, ‘Natural Sounds’, ‘Sounds of Things’, ‘Source Ambiguous Sounds’, and ‘Channel, Environment, Background’.

Low-level visual features include mean perceptual brightness, extracted from the RGB values from each movie frame and mean motion, calculated as the mean of 2,139 motion energy filters from pymoten (***Huth et al., 2012***; ***Naselaris et al., 2011***; ***Nunez-Elizalde et al., 2022***). High-level visual features here include hand-coded labels for the presence of faces and bodies obtained from the EmoCodes dataset (***Camacho et al., 2022***). While many more features could be extracted and included in the models, here we limit features to this number as encoding models from the pilot study with greater numbers of features resulted in overfitting.

### Encoding Models

The hypotheses in this study are tested via encoding models, specifically looking at model performance, unique variance explained, and model weights. Individual subject grayordinate-wise stacked regression (***Lin et al., 2024***) is used, training a separate encoding model for each grayordinate to predict withheld brain activation for each subject. Three different stacked encoding models are employed here: a low- and high-level audio model with only audio features, a low- and high-level visual model with only visual features, and an audio and visual model with all audio and visual features (the audiovisual stacked model is depicted in ***Figure 12***). For each model and subject, the significance of each grayordinate is tested via a null model by repeatedly temporally permuting the order of observations and retraining and testing the models. 1,000 iterations are performed for each subject. FDR correction is applied to account for multiple comparisons for modeling and testing a separate model for each grayordinate across the brain. Model weights and performance are interpreted by pooling data across significant grayordinates within parcels of interest. The encoding models use 5-fold cross-validation, training on 80% of the movie data and testing on the remaining 20% across successive folds. To account for the hemodynamic response, all features are convolved with a canonical double gamma HRF function. Features are z-scored to reduce the effect of scaling on model weights. Accuracy is quantified for each region as the mean coefficient of determination (*R^2^*) and mean noise ceiling normalized *R^2^*. Spearman-Brown corrected split-half noise ceilings are calculated to estimate the upper bound of possible performance for each brain parcel. Unique low-level model *R^2^* is calculated by subtracting the high-level model *R^2^* from the stacked encoding model *R^2^*. For this calculation, when high-level *R^2^* is less than zero, it is clipped to zero, resulting in low-level *R^2^* = stacked *R^2^*. When high-level *R^2^* exceeds stacked *R^2^* it is set to stacked *R^2^*, resulting in low-level *R^2^* = 0. Model weight difference is calculated by subtracting the high-level stacked encoding model weight *α_H_* from the low-level weight *α_L_* (for the audiovisual model, the visual weight *α_V_* is subtracted from the audio weight *α_A_*) resulting in a value [-1,1] where 1 indicates a maximum high-level preference, -1 indicates a maximum low-level preference and 0 indicates no preference between features.

**Figure 11.**
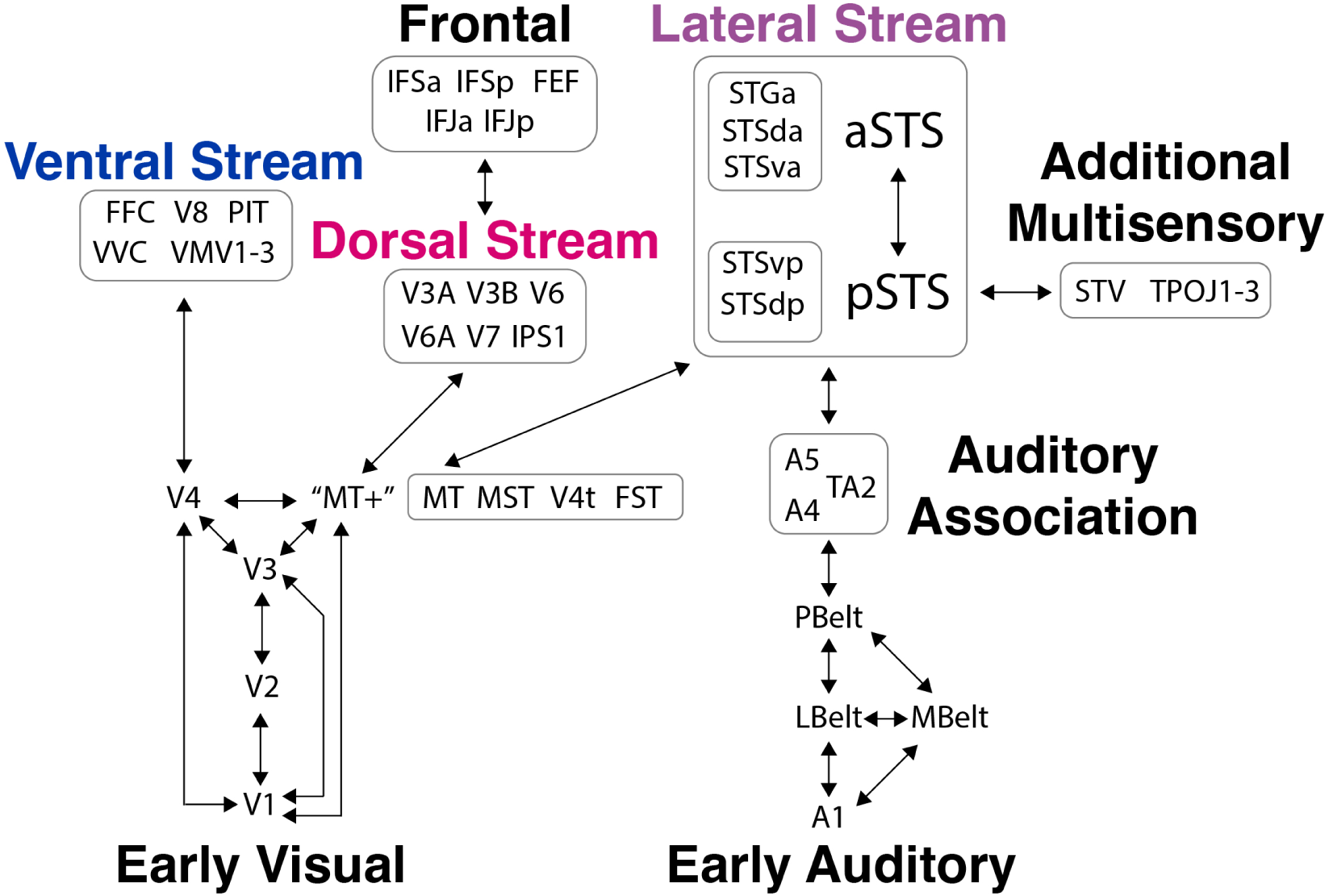
Brain regions of interest with labels selected from the MMP span the visual and auditory systems and the dorsal, ventral, and lateral streams. The schematic shows all perceptual regions of interest grouped by their classification from the MMP and with perceptual streams labeled and some simplified connections illustrated. Note that the actual connectivity between these regions is known to be far more complex, for vision, see ***Felleman and Van Essen*** (1991).

**Figure 12.**
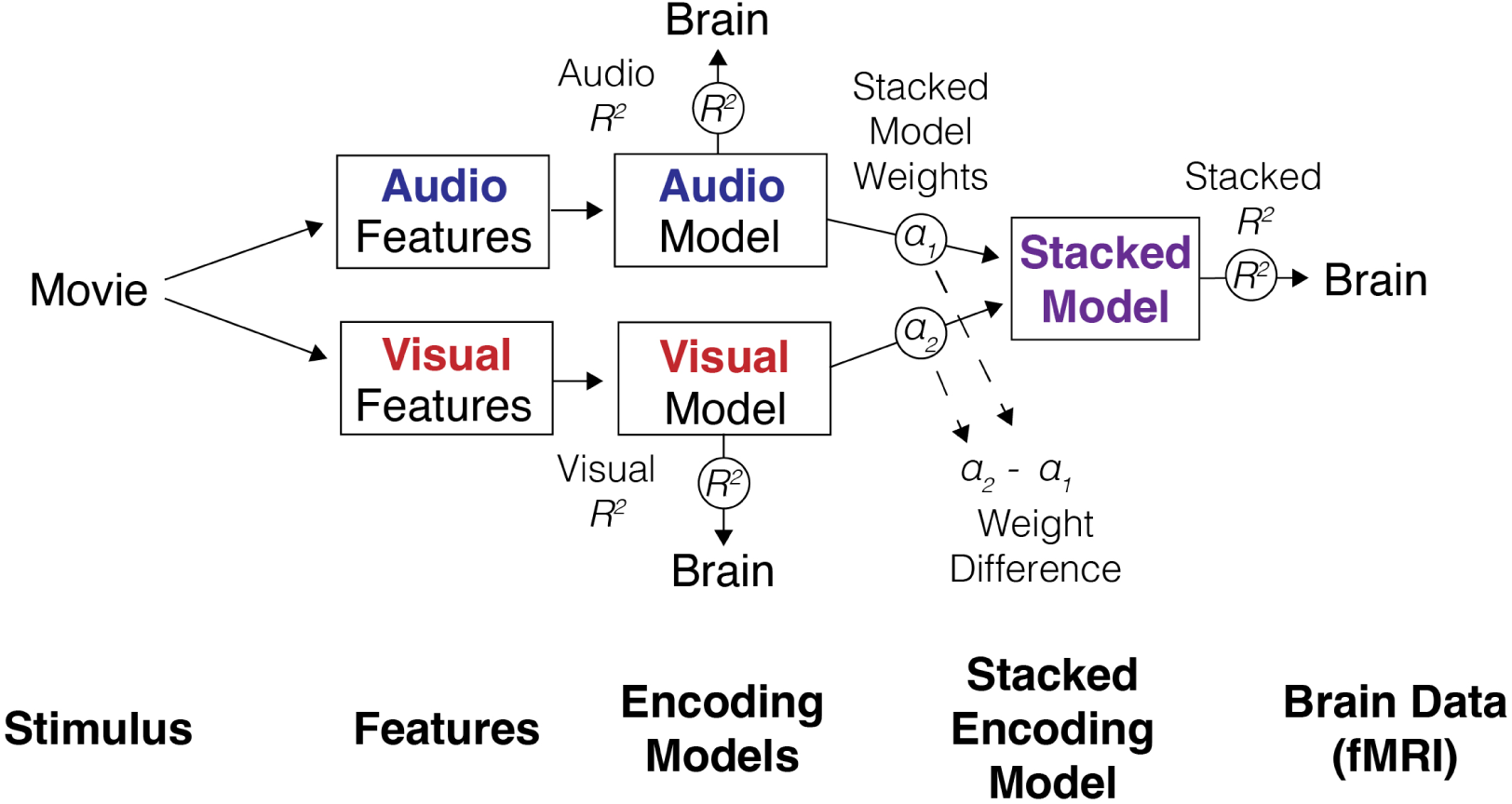
Schematic diagram illustrating the stacked encoding model approach used to relate movie stimulus features to brain activation measured via fMRI in the case of the audio and visual stacked model. From left to right: audio and visual features are extracted from a naturalistic movie stimulus. Each feature set independently informs its own ridge regression encoding model, which predicts brain activation (*R*^2^) on held-out data. The audio and visual ridge regression models are subsequently combined into a stacked regression model, assigning each model a weight (*α*_1_*, α*_2_). The performance of the stacked model is measured by predicting overall brain activation (*R*^2^). The difference between stacked model weights (*α*_2_ − *α*_1_) quantifies the relative perceptual preference toward audio or visual features.

### Statistical Analysis

Linear mixed-effect models are used for significance testing for all hypotheses. While testing the relationship between model performance (*R^2^*) and diagnostic group membership (control vs autism), age, sex, socioeconomic status (SES), and site-level variability are accounted for as covariates. IQ is not included as a covariate in the model due to too many participants with missing data, but a multivariate logistic regression including all covariates and WISC full-scale IQ (the IQ assessment with the largest coverage across the included participants) confirmed that only SRS and sex remained significant independent predictors of diagnostic status. *R^2^*, the coefficient of determination, here represents model fit and serves as the dependent variable. Fixed effects include group membership (binary: 0 or 1), age (continuous, in years), sex (binary: 0 = Female, 1 = Male), and SES (continuous). Site (categorical, across 3 sites) is included as a random intercept to account for unmeasured heterogeneity across sites, allowing the model to adjust for clustering effects. A generalized linear mixed-effects model (GLMM) is fit using a logit link function. The model is implemented in Python using the *pymer4* library (***Jolly, 2018***). Fixed-effect estimates are tested for statistical significance, and the variance of the random effect is assessed to quantify site-level heterogeneity. This same procedure is applied to test the other dependent variables in the low- vs. high-level and audio vs. visual hypotheses: unique explained variance and model weight difference. To test for a relationship with autism severity and sensory symptoms, separate mixed-effects models are employed for each of these metrics, but with the behavioral measures SRS and SSS each included as independent variables of interest in place of diagnostic group membership.

## Acknowledgments

We would like to thank Cat Camacho along with the members of the Senseable Intelligence Group for their insightful discussions.

---

**Figure 2—figure supplement 1.**
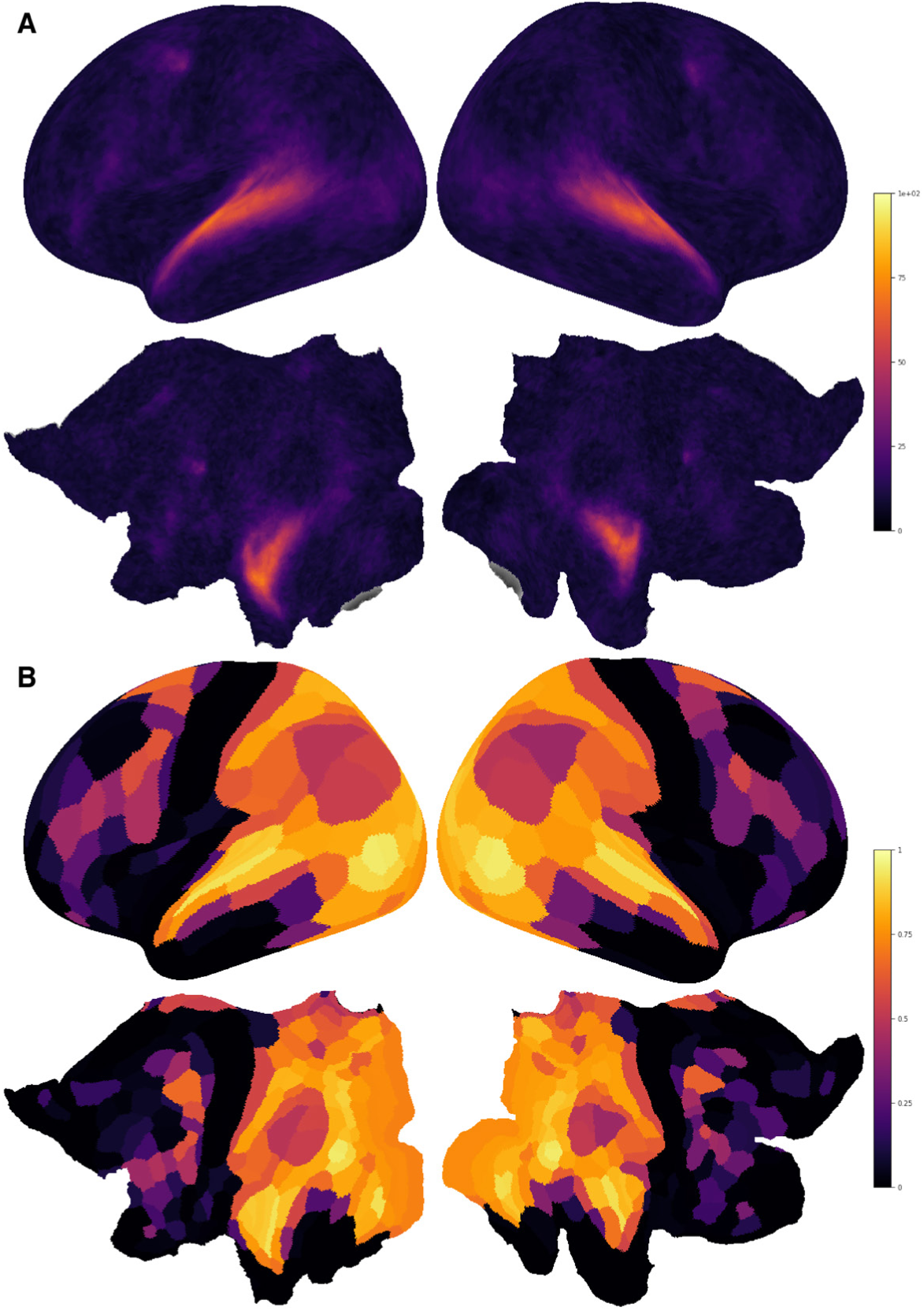
Overall model performance across the cortex. (A) Whole-brain plot of the percentage of subjects with a significant grayordinate at each region. The significance of each grayordinate was tested via a null model by repeatedly temporally permuting the order of observations and retraining and testing the models over 1,000 permutations. (B) The Spearman-Brown corrected split-half noise ceilings for each MMP parcel.

**Figure 2—figure supplement 2.**
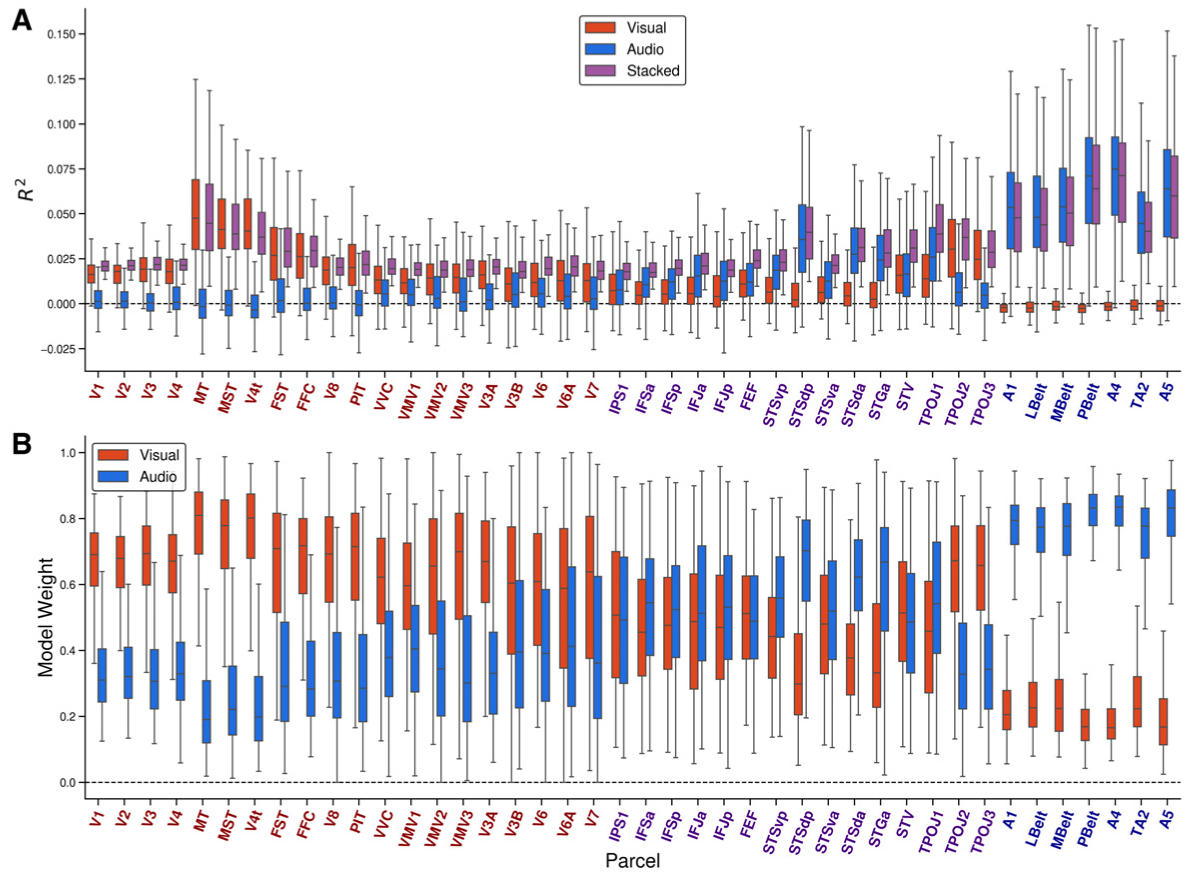
(A) Model performance (*R*^2^) for the visual (red), audio (blue), and stacked (purple) encoding models across perceptual regions of interest for all included participants (both ASD and nonASD). (B) Visual (red) and audio (blue) model weights from the stacked encoding model.

**Figure 2—figure supplement 3.**
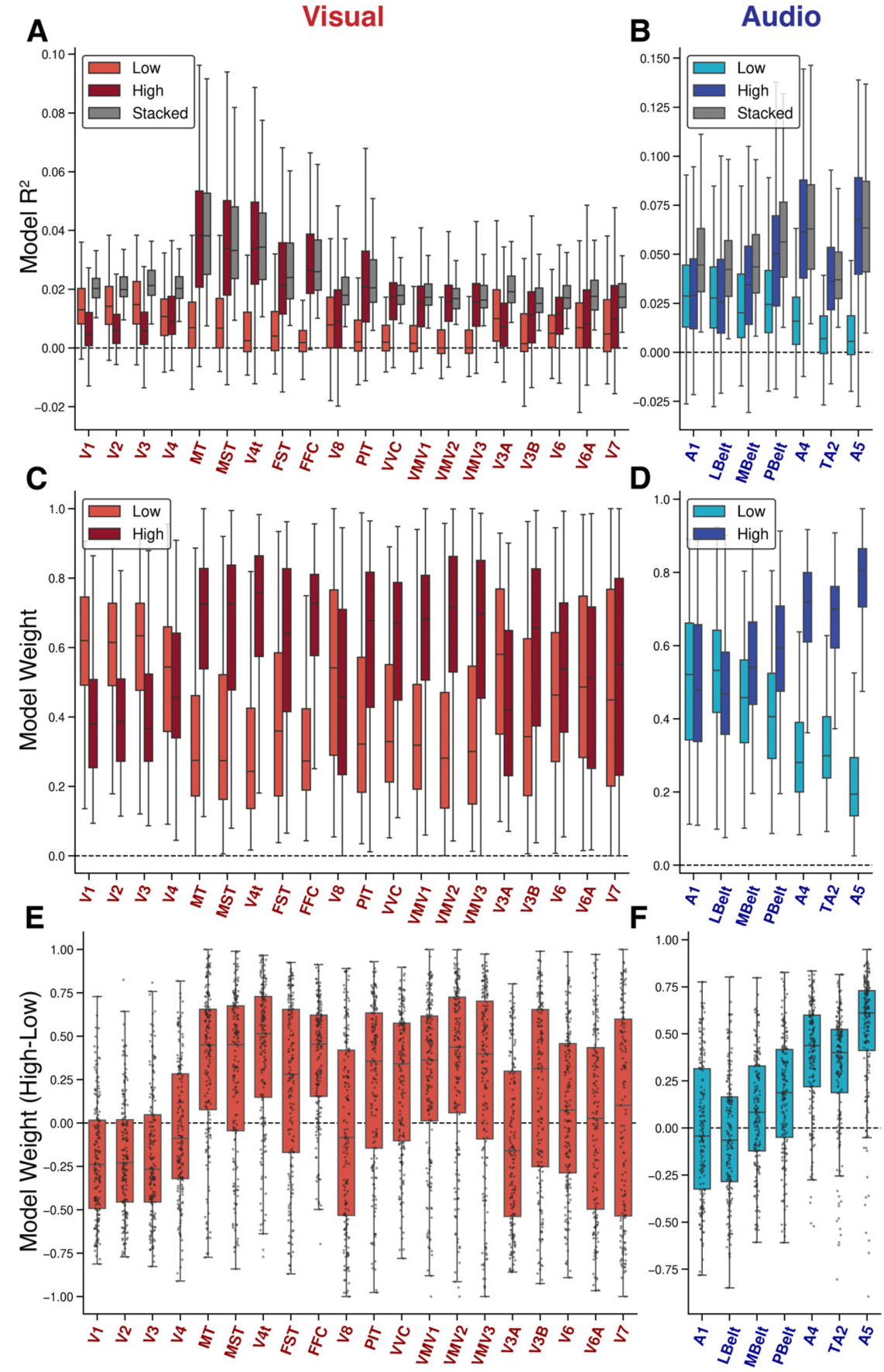
(A) Model performance (R2) for the visual low- (light red) and high-level (dark red) and stacked (grey) encoding models across all nonASD and ASD participants. (A) Corresponding R2 for audio models. (C) Model weights from the stacked visual encoding model. (D) Model weights from the stacked audio encoding model. (E), (F) High- vs. low-level visual and audio perceptual preferences (*W_H_* − *W_L_*) are calculated by taking the difference of high- and low-level weights (shown in B and C). Perceptual preference is in the range of -1 to 1, as the stacked encoding model weights range from 0 to 1. Positive values here indicate a high-level preference, negative values indicate a low-level preference, and values around zero indicate no preference.

**Figure 2—figure supplement 4.**
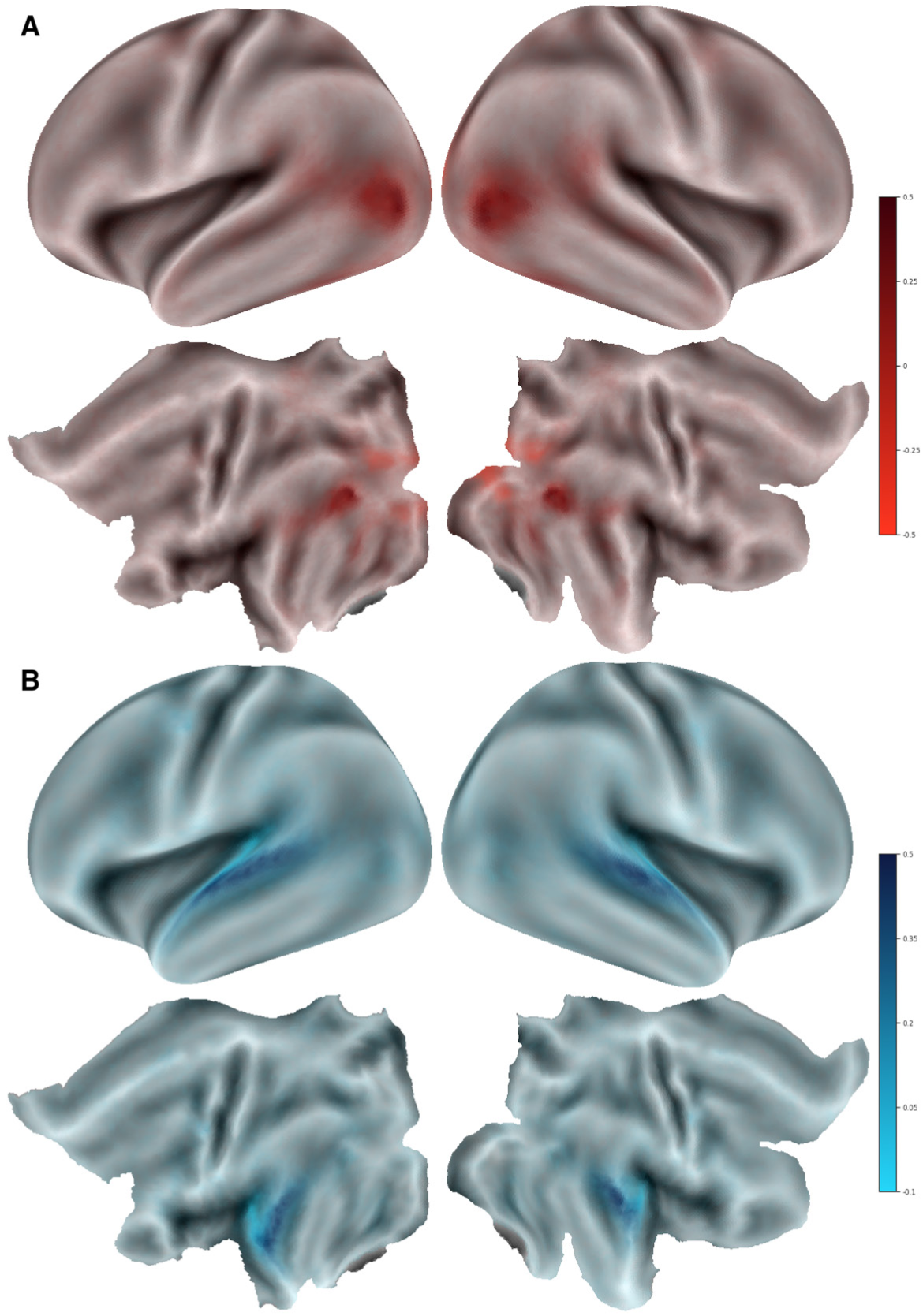
(A) Whole-brain grayordinate-wise plot of mean high- vs. low-level perceptual preference across all participants (ASD and nonASD). (B) The same corresponding plot but from the audio stacked encoding model.

**Figure 3—figure supplement 1.**
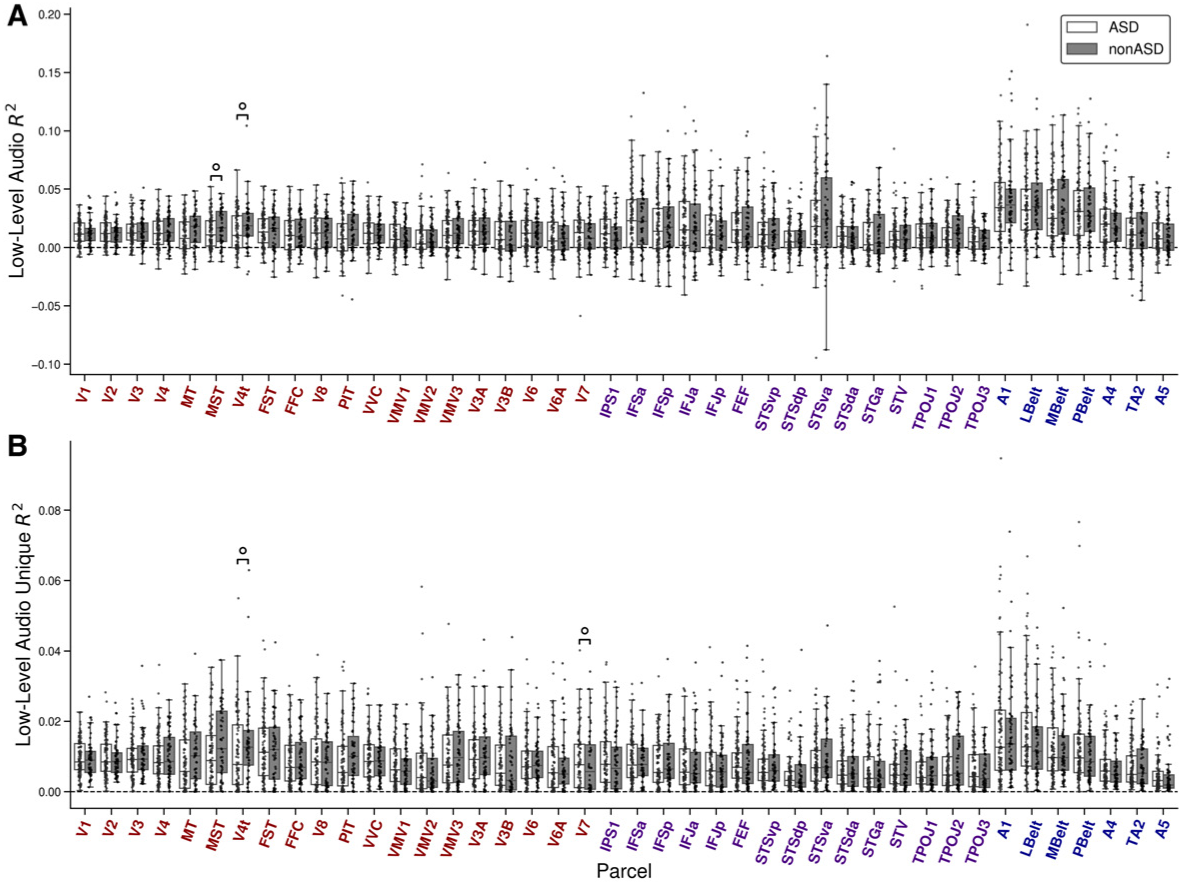
(A) Box plot of low-level audio encoding model explained variance (*R*^2^) for ASD (white) and nonASD (gray) groups across all perceptual ROIs. (B) Box plot of the unique explained variance (*R_u_*^2^). The results correspond to the 40% FD threshold. Boxes annotated with circles indicate an initially significant difference between groups that did not survive FDR correction. Boxes show the quartiles of the dataset, and whiskers show the distribution, with the exception of outliers. Each dot is the mean *R*^2^ from a single subject from all statistically significant grayordinates within each ROI.

**Figure 3—figure supplement 2.**
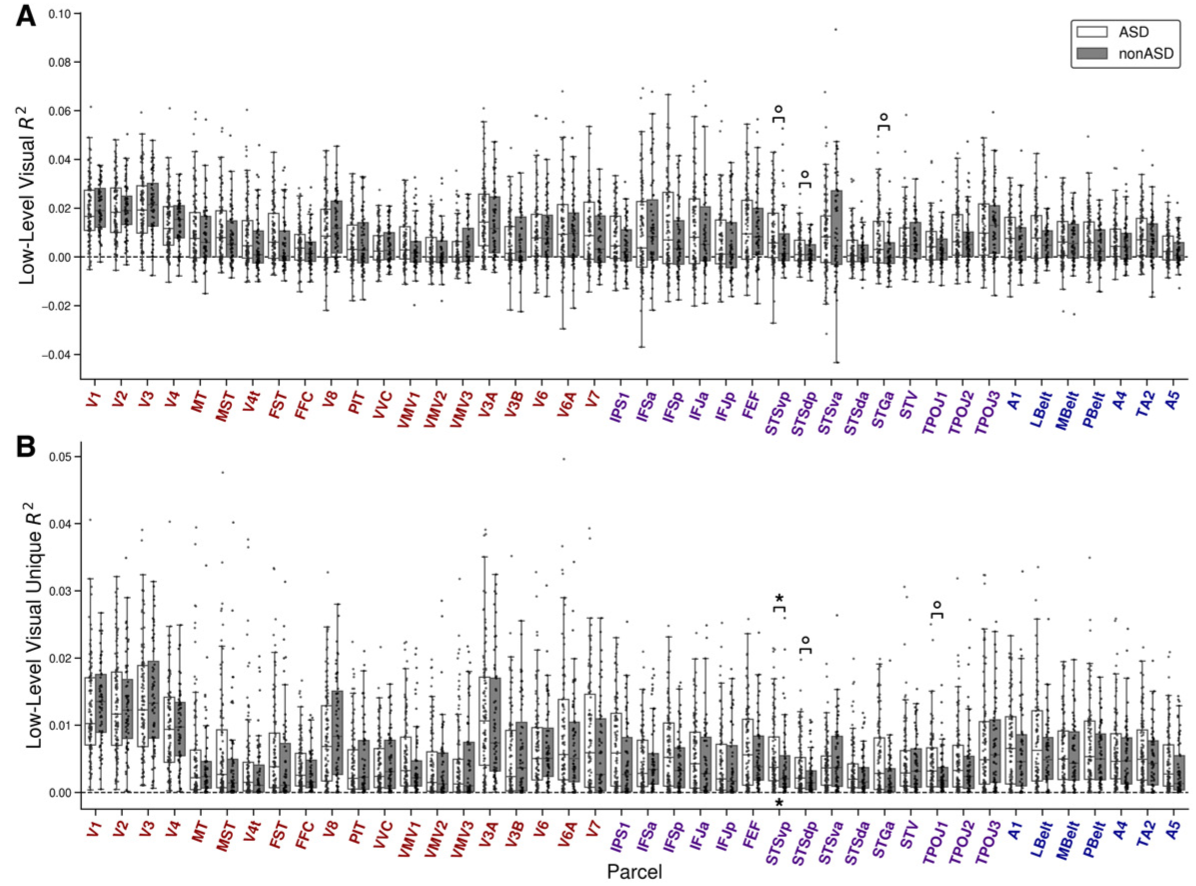
(A) Box plot of low-level visual encoding model *R*^2^ for ASD (white) and nonASD (gray) groups across all perceptual ROIs. (B) Corresponding box plot of *R_u_*^2^. Results correspond to the 40% FD threshold. Boxes annotated with an asterisk indicate a significant group difference (FDR p<0.05), while a circle indicates an initially significant difference between groups that did not survive FDR correction. Boxes show the quartiles of the dataset and whiskers show the distribution with the exception of outliers. Each dot is the mean *R*^2^ from a single subject from all statistically significant grayordinates within each ROI.

**Figure 3—figure supplement 3.**
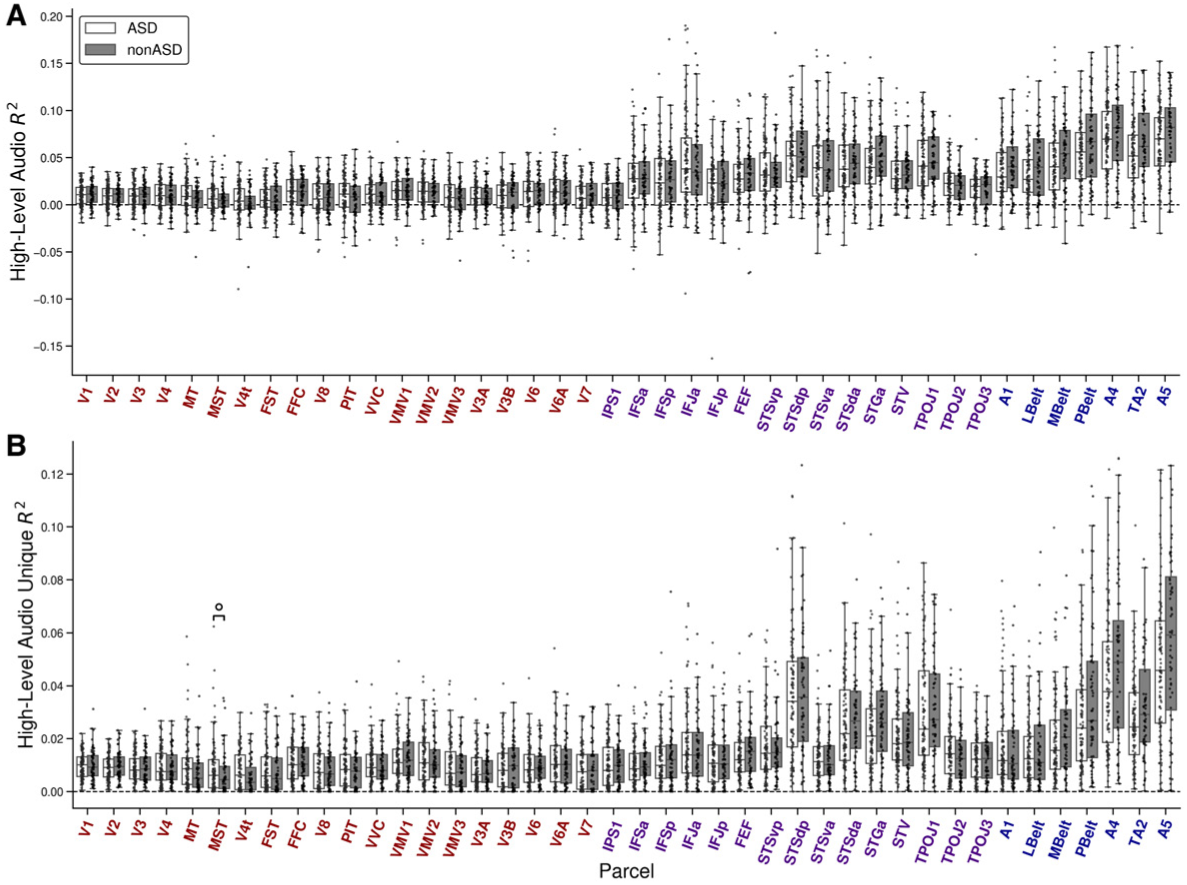
(A) Box plot of high-level audio encoding model *R*^2^ for ASD (white) and nonASD (gray) groups across all perceptual ROIs. (B) Corresponding box plot of *R_u_*^2^. Results correspond to the 40% FD threshold. Boxes annotated with an asterisk indicate a significant group difference (FDR p<0.05), while a circle indicates an initially significant difference between groups that did not survive FDR correction. Boxes show the quartiles of the dataset and whiskers show the distribution with the exception of outliers. Each dot is the mean *R*^2^ from a single subject from all statistically significant grayordinates within each ROI.

**Figure 3—figure supplement 4.**
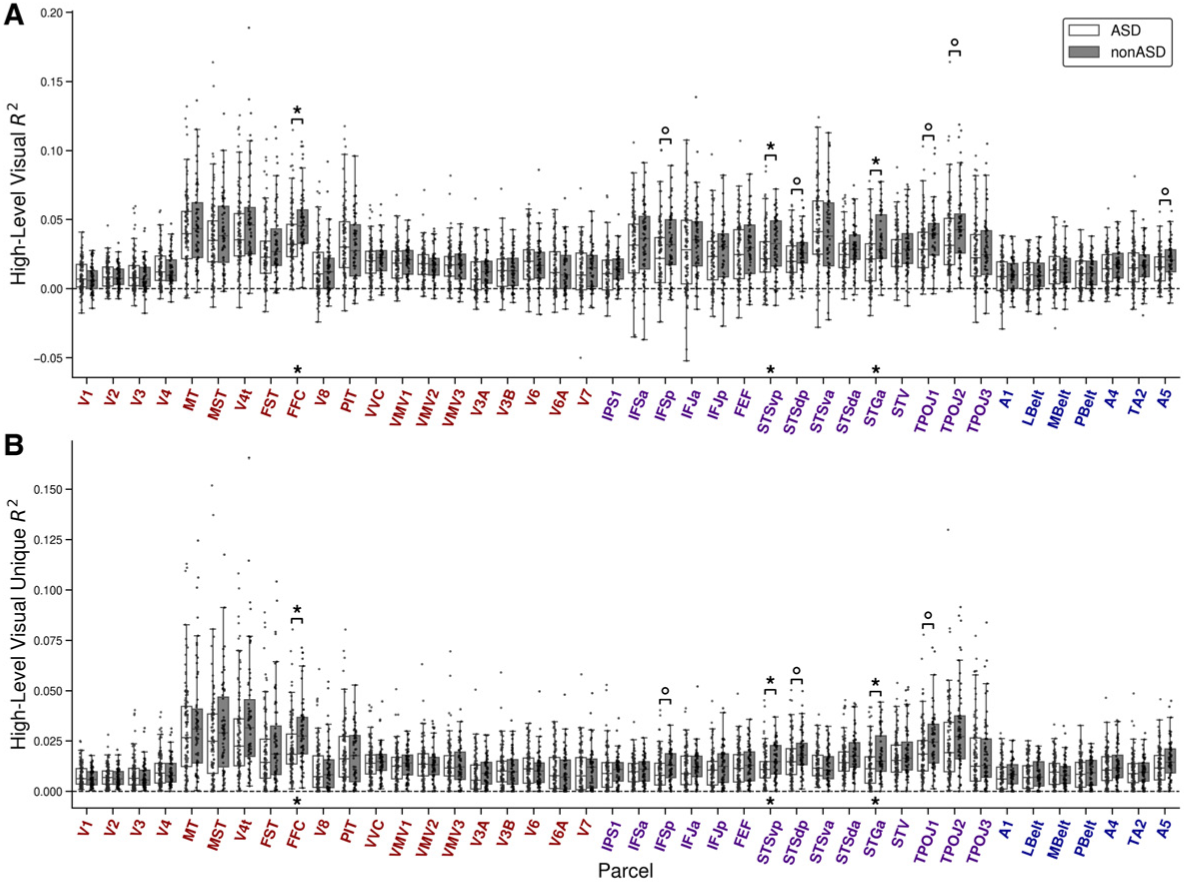
(A) Box plot of high-level visual encoding model *R*^2^ for ASD (white) and nonASD (gray) groups across all perceptual ROIs. (B) Corresponding box plot of *R_u_*^2^. Results correspond to the 40% FD threshold. Boxes annotated with an asterisk indicate a significant group difference (FDR p<0.05), while a circle indicates an initially significant difference between groups that did not survive FDR correction. Boxes show the quartiles of the dataset and whiskers show the distribution with the exception of outliers. Each dot is the mean *R*^2^ from a single subject from all statistically significant grayordinates within each ROI.

**Figure 7—figure supplement 1.**
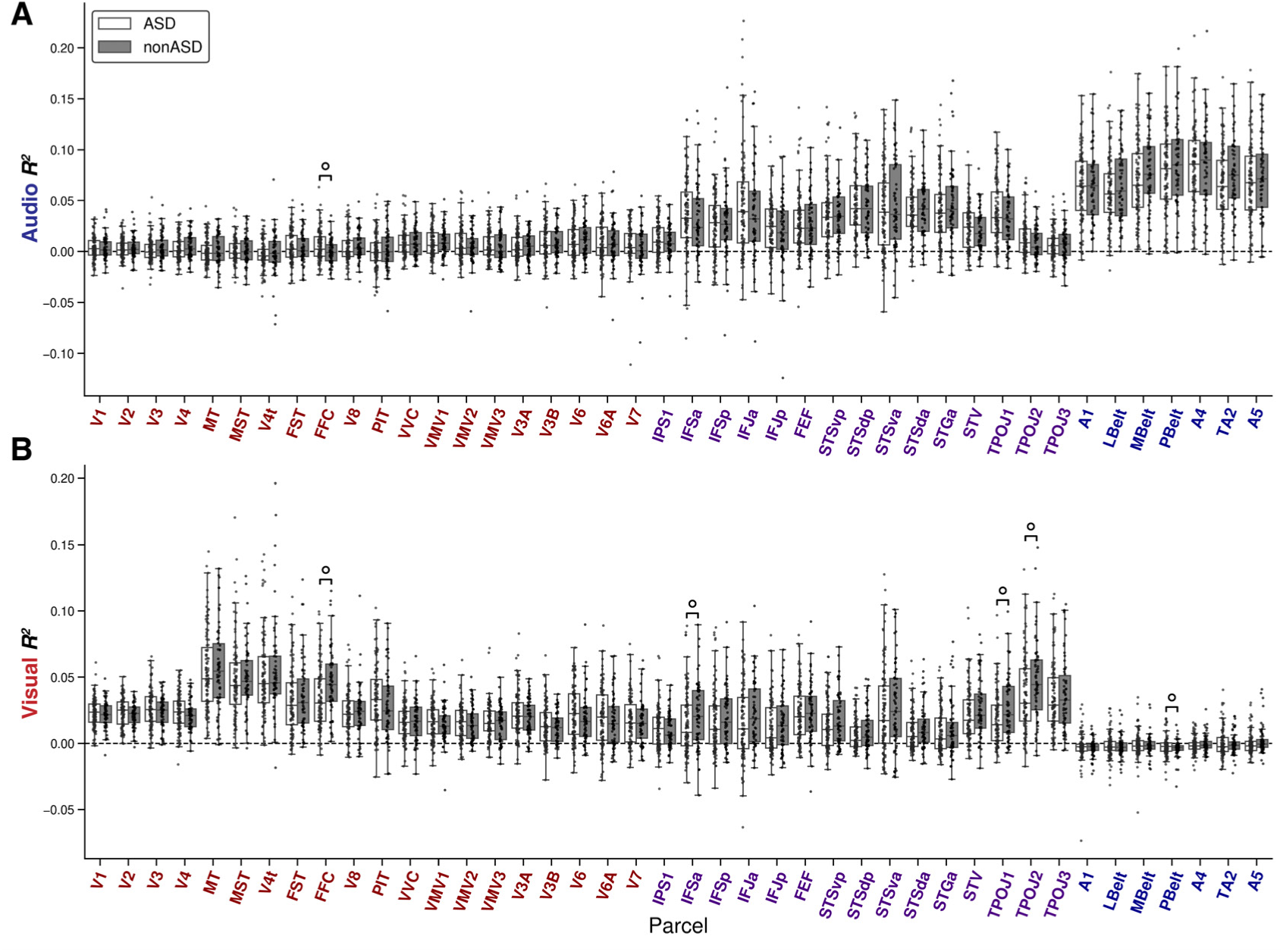
(A) Box plot of audio encoding model *R_u_*^2^ for ASD (white) and nonASD (gray) groups across all perceptual ROIs. (B) The same but for visual *R_u_*^2^. Results correspond with the 40% FD threshold. Boxes annotated with circles indicate a difference between groups that did not survive FDR correction (uncorrected p<0.05, FDR q>0.05). Boxes show the quartiles of the dataset and whiskers show the distribution with the exception of outliers. Each dot is the mean value from a single subject from all statistically significant grayordinates within each ROI.

**Figure 7—figure supplement 2.**
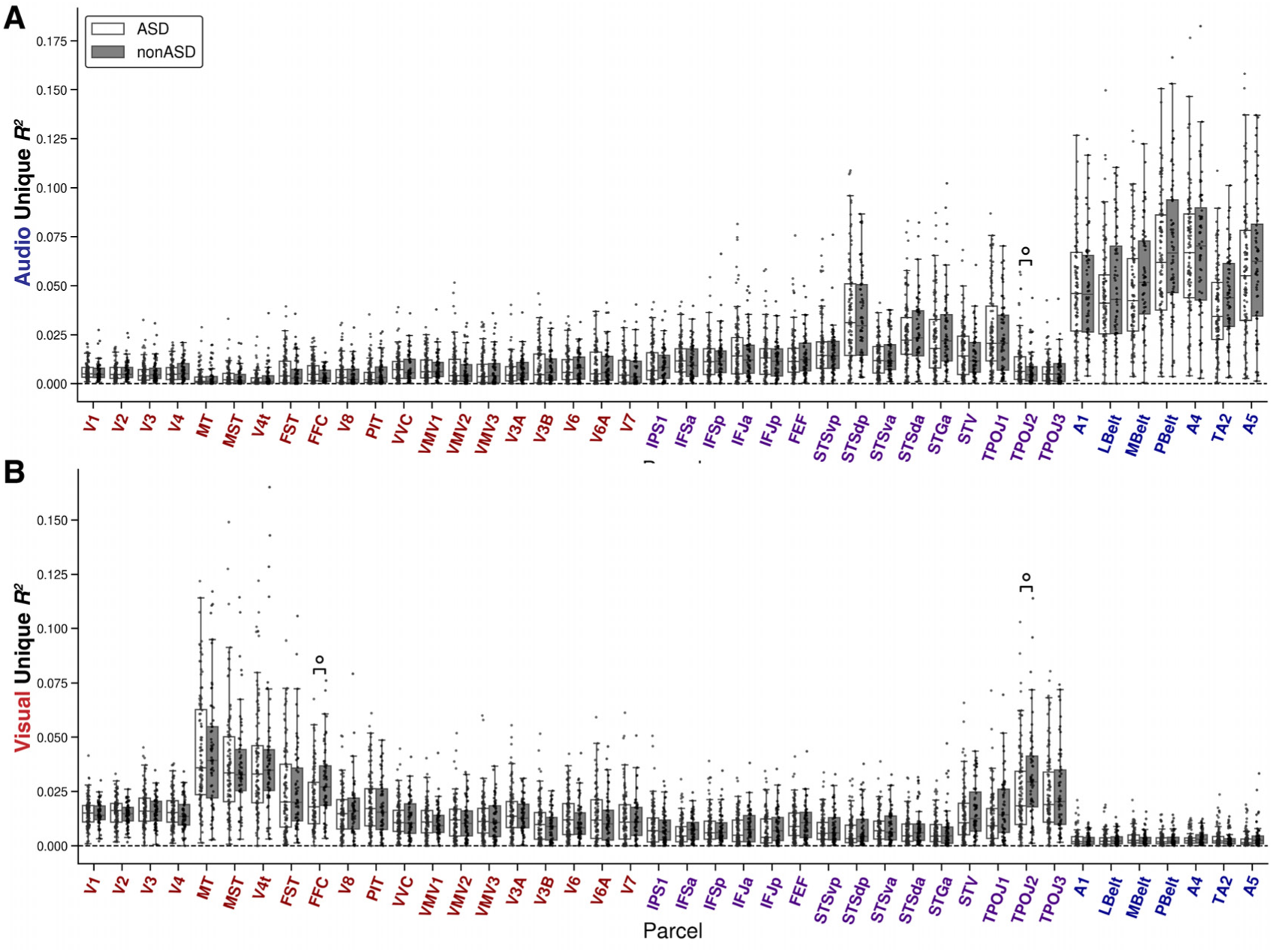
(A) Box plot of audio encoding model *R*^2^ for ASD (white) and nonASD (gray) groups across all perceptual ROIs. (B) The same but for visual *R*^2^. These results correspond with the 40% FD threshold. Boxes annotated with circles indicate a difference between groups that did not survive FDR correction (uncorrected p<0.05, FDR q>0.05). Boxes show the quartiles of the dataset and whiskers show the distribution with the exception of outliers. Each dot is the mean value from a single subject from all statistically significant grayordinates within each ROI.

**Figure 7—figure supplement 3.**
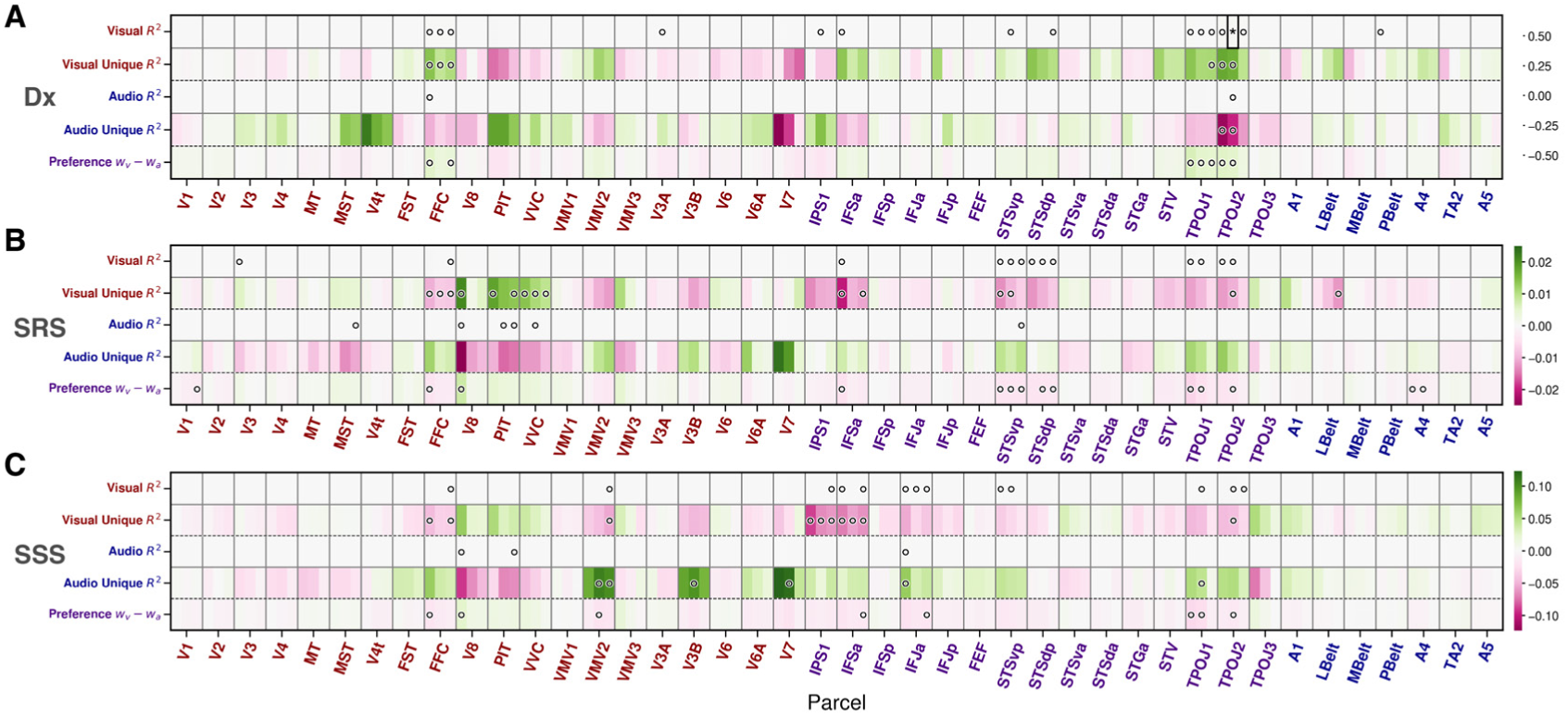
Heatmaps of fixed-effect coefficients for (A) diagnostic group (Dx: nonASD vs. ASD), (B) SRS Total T-Score and (C) sensory subset score (SSS), across 40%, 60% and 80% FD thresholds (left to right within each cortical parcel column). Within each of the three horizontal panels, rows denote encoding model-derived metrics (visual *R*^2^ and *R_u_*^2^, audio *R*^2^ and *R_u_*^2^, and their preference index (*W_V_* − *W_A_*) and columns denote Glasser MMP ROIs ordered left to right from early visual areas through association cortices followed by auditory areas. Color indicates the magnitude and sign of the coefficient (pink=negative effect with ASD>nonASD; green=positive effect with ASD<nonASD). Asterisks mark FDR-corrected significance at *q <* 0.05; open circles mark uncorrected *p <* 0.05. The visual modality encoding models are labeled in red and the auditory in blue.

**Figure 8—figure supplement 1.**
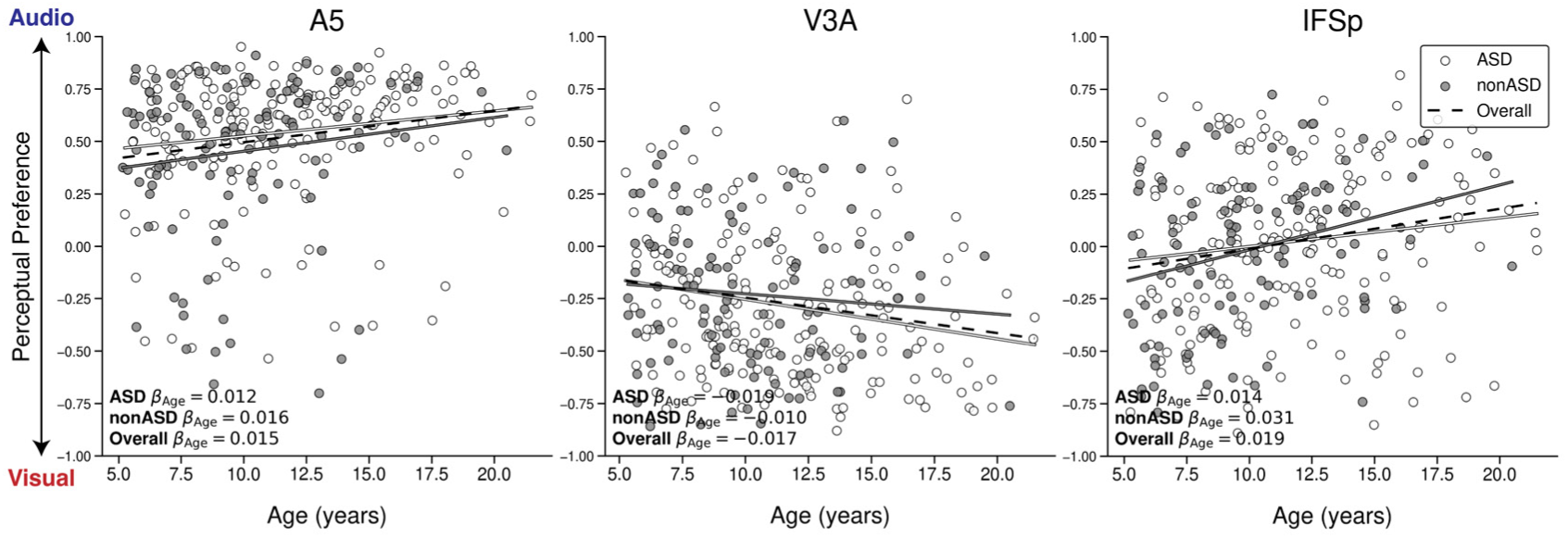
Scatter plots showing the relationship between age and perceptual preference in three example cortical regions (A5, V3A, and IFSp) where significant effects were observed across all participants. Perceptual preference values above zero indicate an auditory preference, and values below zero indicate a visual preference. Although no significant age-by-diagnosis interactions are displayed here, autistic (ASD; white) and non-autistic (nonASD; gray) groups are colored separately for clarity. Lines show model fits for each group colored correspondingly, with estimated age-related slopes (*β_Age_*) reported for ASD, nonASD, and the overall sample (black dashed line).

**Figure 8—figure supplement 2.**
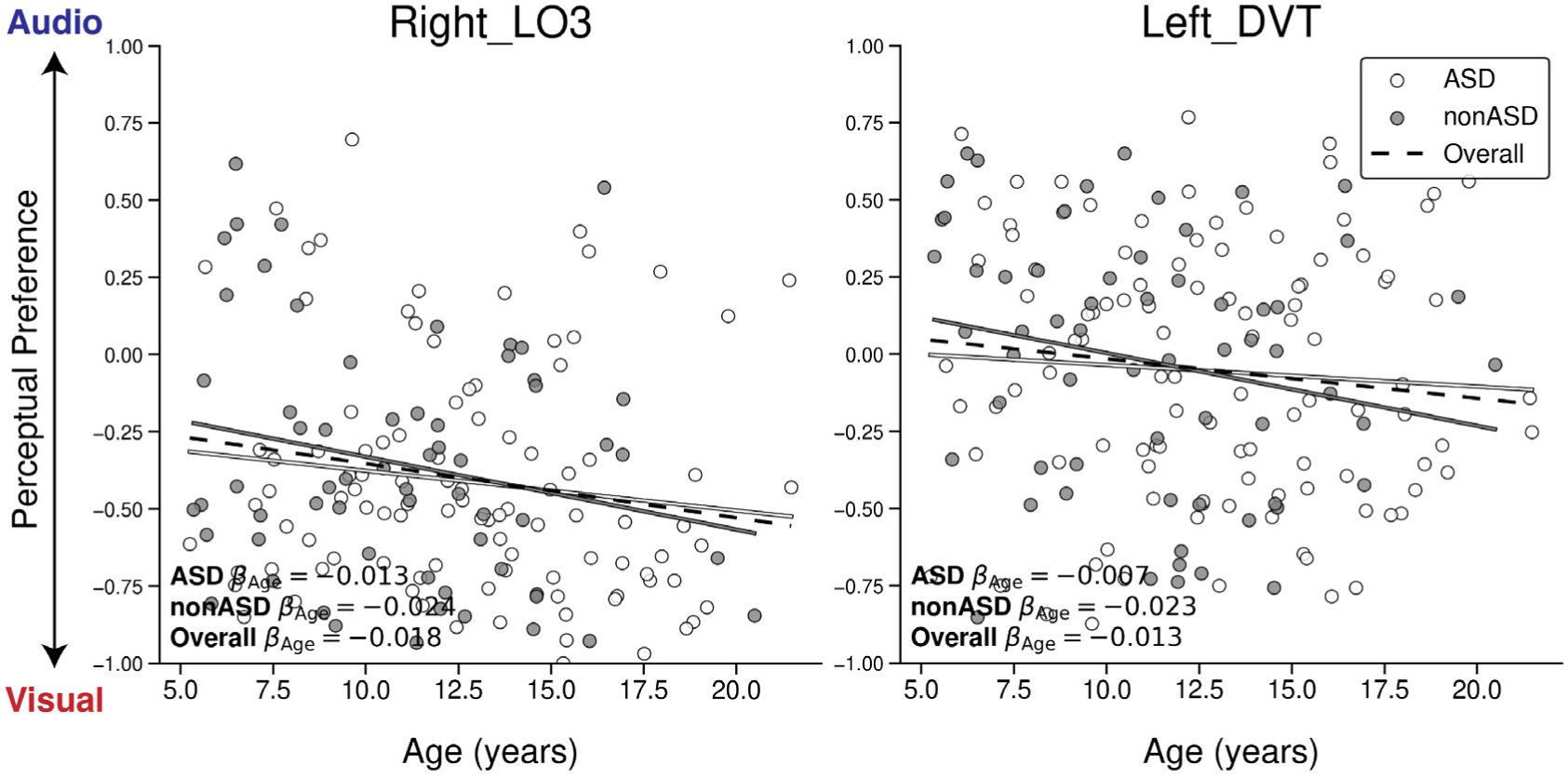
Scatter plots showing the relationship between age and perceptual preference in two example lateralized cortical regions, Right LO3 (a visual region between early visual cortex and MT+) and Left DVT (Dorsal Visual Transitional area; a region located on the posterior bank of the parieto-occipital sulcus), where significant effects were observed across all participants at the whole-brain level. Perceptual preference values above zero indicate an auditory preference, and values below zero indicate a visual preference. Although no significant age-by-diagnosis interactions are displayed here, autistic (ASD; white) and non-autistic (nonASD; gray) groups are colored separately for clarity. Lines show model fits for each group, with estimated age-related slopes (*β_Age_*) reported for ASD, nonASD, and the overall sample (black dashed line).

**Figure 8—figure supplement 3.**
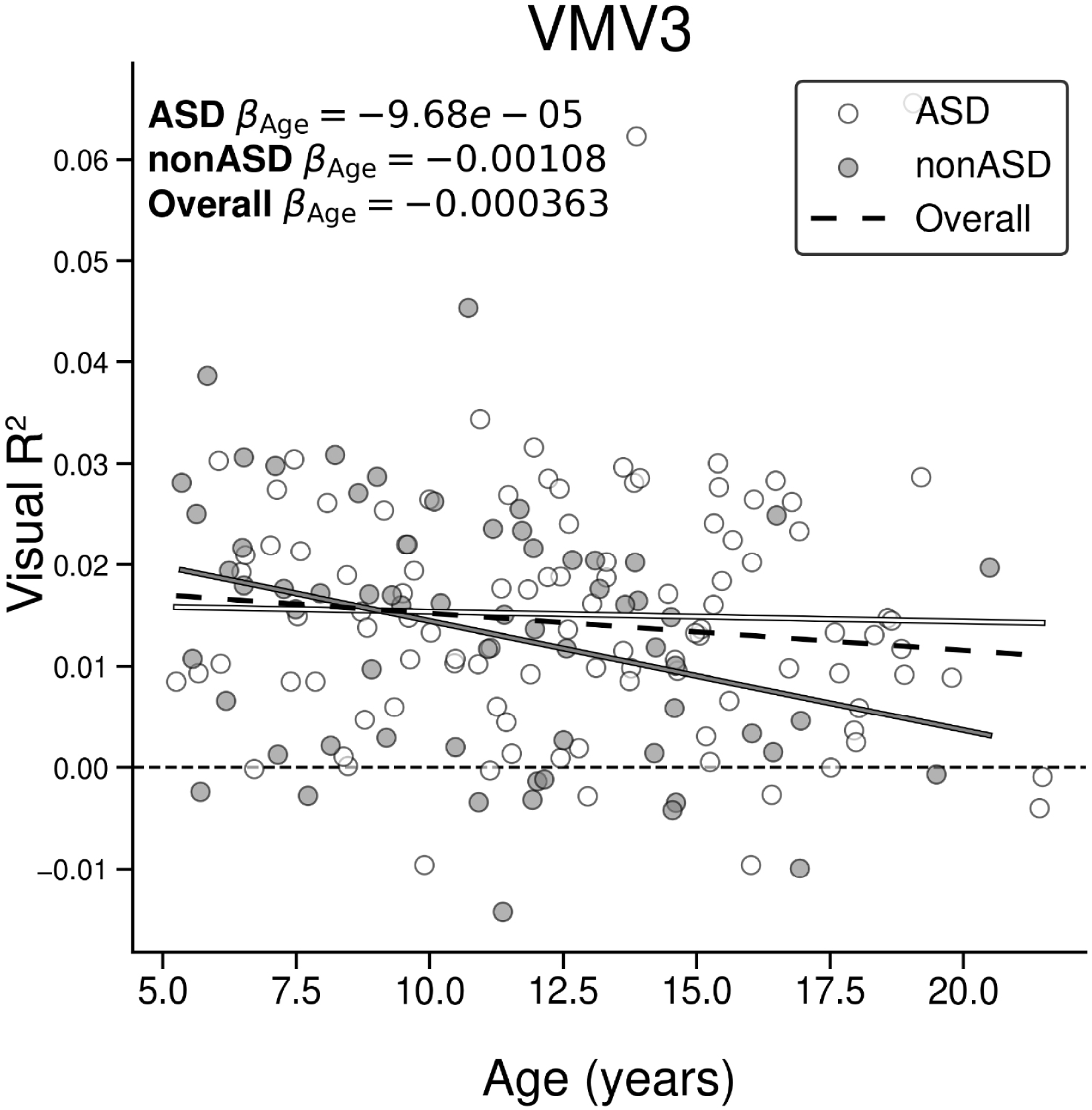
Scatter plot showing the relationship between age and visual *R*^2^ in perceptual region VMV3 where a significant age:diagnosis interaction was observed. Autistic (ASD; white) and non-autistic (nonASD; gray) diagnostic groups are displayed. Lines show model fits for each group, with estimated age-related slopes (*β_Age_*) reported for ASD, nonASD, and the overall sample (black dashed line).

## References

Abrams DA, Lynch CJ, Cheng KM, Phillips J, Supekar K, Ryali S, Uddin LQ, Menon V. Underconnectivity between voice-selective cortex and reward circuitry in children with autism. Proceedings of the National Academy of Sciences. 2013; 110(29):12060–12065.

Alaerts K, Nayar K, Kelly C, Raithel J, Milham MP, Di Martino A. Age-related changes in intrinsic function of the superior temporal sulcus in autism spectrum disorders. Social cognitive and affective neuroscience. 2015; 10(10):1413–1423.

Alaerts K, Woolley DG, Steyaert J, Di Martino A, Swinnen SP, Wenderoth N. Underconnectivity of the superior temporal sulcus predicts emotion recognition deficits in autism. Social cognitive and affective neuroscience. 2014; 9(10):1589–1600.

Alexander LM, Escalera J, Ai L, Andreotti C, Febre K, Mangone A, Vega-Potler N, Langer N, Alexander A, Kovacs M, Litke S, O’Hagan B, Andersen J, Bronstein B, Bui A, Bushey M, Butler H, Castagna V, Camacho N, Chan E, et al. An open resource for transdiagnostic research in pediatric mental health and learning disorders. Sci Data. 2017 Dec; 4:170181.

Balasco L, Provenzano G, Bozzi Y. Sensory abnormalities in autism spectrum disorders: a focus on the tactile domain, from genetic mouse models to the clinic. Frontiers in psychiatry. 2020; 10:1016.

Ben-Sasson A, Cermak SA, Orsmond GI, Tager-Flusberg H, Carter AS, Kadlec MB, Dunn W. Extreme sensory modulation behaviors in toddlers with autism spectrum disorders. The American Journal of Occupational Therapy. 2007; 61(5):584–592.

Bennetto L, Kuschner ES, Hyman SL. Olfaction and taste processing in autism. Biological psychiatry. 2007; 62(9):1015–1021.

Bolton TAW, Freitas LGA, Jochaut D, Giraud AL, Van De Ville D. Neural responses in autism during movie watching: Inter-individual response variability co-varies with symptomatology. Neuroimage. 2020 Aug; 216:116571.

Bolton TAW, Jochaut D, Giraud AL, Van De Ville D. Brain dynamics in ASD during movie-watching show idiosyncratic functional integration and segregation. Hum Brain Mapp. 2018 Jun; 39(6):2391–2404.

Brock J. Alternative Bayesian accounts of autistic perception: comment on Pellicano and Burr. Trends Cogn Sci. 2012 Dec; 16(12):573–4; author reply 574–5.

Burr D, Gori M. Multisensory Integration Develops Late in Humans. In: Murray MM, Wallace MT, editors. The Neural Bases of Multisensory Processes Frontiers in Neuroscience, Boca Raton, FL: CRC Press/Taylor & Francis; 2012.https://www.ncbi.nlm.nih.gov/books/NBK92864/.

Byrge L, Dubois J, Tyszka JM, Adolphs R, Kennedy DP. Idiosyncratic brain activation patterns are associated with poor social comprehension in autism. J Neurosci. 2015 Apr; 35(14):5837–5850.

Bölte S, Poustka F, Constantino JN. Assessing autistic traits: cross-cultural validation of the social responsiveness scale (SRS). Autism Res. 2008 Dec; 1(6):354–363.

Calvert GA, Campbell R, Brammer MJ. Evidence from functional magnetic resonance imaging of crossmodal binding in the human heteromodal cortex. Curr Biol. 2000 Jun; 10(11):649–657.

Calvert GA, Hansen PC, Iversen SD, Brammer MJ. Detection of audio-visual integration sites in humans by application of electrophysiological criteria to the BOLD effect. Neuroimage. 2001; 14(2):427–438.

Camacho MC, Nielsen AN, Balser D, Furtado E, Steinberger DC, Fruchtman L, Culver JP, Sylvester CM, Barch DM. Large-scale encoding of emotion concepts becomes increasingly similar between individuals from childhood to adolescence. Nat Neurosci. 2023 Jun; .

Camacho MC, Williams EM, Balser D, Kamojjala R, Sekar N, Steinberger D, Yarlagadda S, Perlman SB, Barch DM. EmoCodes: A standardized coding system for Socio-emotional content in complex video stimuli. Affect Sci. 2022 Mar; 3(1):168–181.

Cary E, Pacheco D, Kaplan-Kahn E, McKernan E, Matsuba E, Prieve B, Russo N. Brain signatures of early and late neural measures of auditory habituation and discrimination in autism and their relationship to autistic traits and sensory overresponsivity. Journal of Autism and Developmental Disorders. 2024; 54(4):1344–1360.

Cascio CJ, Woynaroski T, Baranek GT, Wallace MT. Toward an interdisciplinary approach to understanding sensory function in autism spectrum disorder. Autism Res. 2016 Sep; 9(9):920–925.

Cerliani L, Mennes M, Thomas RM, Di Martino A, Thioux M, Keysers C. Increased functional connectivity between subcortical and cortical resting-state networks in autism spectrum disorder. JAMA psychiatry. 2015; 72(8):767–777.

Chamak B, Bonniau B, Jaunay E, Cohen D. What can we learn about autism from autistic persons? Psychotherapy and psychosomatics. 2008; 77(5):271–279.

Chen YJ, Duku E, Georgiades S. Rethinking autism intervention science: A dynamic perspective. Frontiers in psychiatry. 2022; 13:827406.

Cohen SS, Tottenham N, Baldassano C. Developmental changes in story-evoked responses in the neocortex and hippocampus. Elife. 2022 Jul; 11.

Colavita FB. Human sensory dominance. Perception & Psychophysics. 1974 Oct; 16(2):409–412.

Constantino JN. Social Responsiveness Scale (SRS). Western Psychological Services; 2007.

Constantino JN, Davis SA, Todd RD, Schindler MK, Gross MM, Brophy SL, Metzger LM, Shoushtari CS, Splinter R, Reich W. Validation of a brief quantitative measure of autistic traits: comparison of the social responsiveness scale with the autism diagnostic interview-revised. J Autism Dev Disord. 2003 Aug; 33(4):427–433.

Courchesne E, Pierce K. Why the frontal cortex in autism might be talking only to itself: local over-connectivity but long-distance disconnection. Curr Opin Neurobiol. 2005 Apr; 15(2):225–230.

Van de Cruys S, Evers K, Van der Hallen R, Van Eylen L, Boets B, de Wit L, Wagemans J. Precise minds in uncertain worlds: predictive coding in autism. Psychol Rev. 2014 Oct; 121(4):649–675.

Dakin S, Frith U. Vagaries of visual perception in autism. Neuron. 2005 Nov; 48(3):497–507.

Damarla SR, Keller TA, Kana RK, Cherkassky VL, Williams DL, Minshew NJ, Just MA. Cortical underconnectivity coupled with preserved visuospatial cognition in autism: Evidence from an fMRI study of an embedded figures task. Autism Res. 2010 Oct; 3(5):273–279.

Di X, Xu T, Uddin LQ, Biswal BB. Individual differences in time-varying and stationary brain connectivity during movie watching from childhood to early adulthood: Age, sex, and behavioral associations. Dev Cogn Neurosci. 2023 Jul; 63:101280.

Di Martino A, Yan CG, Li Q, Denio E, Castellanos FX, Alaerts K, Anderson JS, Assaf M, Bookheimer SY, Dapretto M, Deen B, Delmonte S, Dinstein I, Ertl-Wagner B, Fair DA, Gallagher L, Kennedy DP, Keown CL, Keysers C, Lainhart JE, et al. The autism brain imaging data exchange: towards a large-scale evaluation of the intrinsic brain architecture in autism. Mol Psychiatry. 2014 Jun; 19(6):659–667.

Diedrichsen J, Kriegeskorte N. Representational models: A common framework for understanding encoding, pattern-component, and representational-similarity analysis. PLoS Comput Biol. 2017 Apr; 13(4):e1005508.

Dinstein I, Heeger DJ, Lorenzi L, Minshew NJ, Malach R, Behrmann M. Unreliable evoked responses in autism. Neuron. 2012 Sep; 75(6):981–991.

DuBois D, Ameis SH, Lai MC, Casanova MF, Desarkar P. Interoception in autism spectrum disorder: A review. International journal of developmental neuroscience. 2016; 52:104–111.

Elyounssi S, Kunitoki K, Clauss JA, Laurent E, Kane KA, Hughes DE, Hopkinson CE, Bazer O, Sussman RF, Doyle AE, et al. Addressing artifactual bias in large, automated MRI analyses of brain development. Nature Neuroscience. 2025; p. 1–10.

Esteban O, Markiewicz CJ, Blair RW, Moodie CA, Isik AI, Erramuzpe A, Kent JD, Goncalves M, DuPre E, Snyder M, Oya H, Ghosh SS, Wright J, Durnez J, Poldrack RA, Gorgolewski KJ. fMRIPrep: a robust preprocessing pipeline for functional MRI. Nat Methods. 2019 Jan; 16(1):111–116.

Falck-Ytter T, Bussu G. The sensory-first account of autism. Neurosci Biobehav Rev. 2023 Oct; 153:105405.

Federici A, Parma V, Vicovaro M, Radassao L, Casartelli L, Ronconi L. Anomalous perception of biological motion in autism: a conceptual review and meta-analysis. Scientific Reports. 2020; 10(1):4576.

Feldman JI, Conrad JG, Kuang W, Tu A, Liu Y, Simon DM, Wallace MT, Woynaroski TG. Relations between the McGurk effect, social and communication skill, and autistic features in children with and without autism. J Autism Dev Disord. 2022 May; 52(5):1920–1928.

Feldman JI, Dunham K, Cassidy M, Wallace MT, Liu Y, Woynaroski TG. Audiovisual multisensory integration in individuals with autism spectrum disorder: A systematic review and meta-analysis. Neurosci Biobehav Rev. 2018 Dec; 95:220–234.

Felleman DJ, Van Essen DC. Distributed hierarchical processing in the primate cerebral cortex. Cereb Cortex. 1991; 1(1):1–47.

Floris DL, Llera A, Zabihi M, Moessnang C, Jones EJ, Mason L, Haartsen R, Holz NE, Mei T, Elleaume C, et al. A multimodal neural signature of face processing in autism within the fusiform gyrus. Nature Mental Health. 2025; 3(1):31–45.

Friston K. A theory of cortical responses. Philosophical transactions of the Royal Society B: Biological sciences. 2005; 360(1456):815–836.

Frith U. Autism: Explaining the Enigma. Oxford, UK: Blackwell; 1989.

Gemmeke JF, Ellis DPW, Freedman D, Jansen A, Lawrence W, Moore RC, Plakal M, Ritter M. Audio Set: An ontology and human-labeled dataset for audio events. In: 2017 IEEE International Conference on Acoustics, Speech and Signal Processing (ICASSP) IEEE; 2017. p. 776–780.

Georgiades S, Bishop SL, Frazier T, Editorial Perspective: Longitudinal research in autism–introducing the concept of ‘chronogeneity’. Wiley Online Library; 2017.

Ghazanfar AA, Schroeder CE. Is neocortex essentially multisensory? Trends Cogn Sci. 2006 Jun; 10(6):278–285.

Glasser MF, Coalson TS, Robinson EC, Hacker CD, Harwell J, Yacoub E, Ugurbil K, Andersson J, Beckmann CF, Jenkinson M, Smith SM, Van Essen DC. A multi-modal parcellation of human cerebral cortex. Nature. 2016 Aug; 536(7615):171–178.

Guo X, He C, Duan X, Han S, Xiao J, Chen H. Aberrant functional connectivity dynamics of superior temporal sulcus and its associations with GABA genes expression in autism. In: Proceedings of the 3rd International Conference on Medical and Health Informatics; 2019. p. 21–25.

Hadjikhani N, Joseph RM, Snyder J, Tager-Flusberg H. Abnormal activation of the social brain during face perception in autism. Hum Brain Mapp. 2007 May; 28(5):441–449.

Han S, Tai C, Westenbroek RE, Yu FH, Cheah CS, Potter GB, Rubenstein JL, Scheuer T, De La Iglesia HO, Catterall WA. Autistic-like behaviour in Scn1a+/- mice and rescue by enhanced GABA-mediated neurotransmission. Nature. 2012; 489(7416):385–390.

Happé FG. Studying weak central coherence at low levels: children with autism do not succumb to visual illusions. A research note. Journal of child psychology and psychiatry. 1996; 37(7):873–877.

Happé F, Ronald A, Plomin R. Time to give up on a single explanation for autism. Nat Neurosci. 2006 Oct; 9(10):1218–1220.

Hasson U, Avidan G, Gelbard H, Vallines I, Harel M, Minshew N, Behrmann M. Shared and idiosyncratic cortical activation patterns in autism revealed under continuous real-life viewing conditions. Autism Res. 2009 Aug; 2(4):220–231.

Haxby JV, Connolly AC, Guntupalli JS. Decoding neural representational spaces using multivariate pattern analysis. Annu Rev Neurosci. 2014 Jun; 37:435–456.

Haxby JV, Guntupalli JS, Connolly AC, Halchenko YO, Conroy BR, Gobbini MI, Hanke M, Ramadge PJ. A common, high-dimensional model of the representational space in human ventral temporal cortex. Neuron. 2011 Oct; 72(2):404–416.

Herbert MR, Harris GJ, Adrien KT, Ziegler DA, Makris N, Kennedy DN, Lange NT, Chabris CF, Bakardjiev A, Hodgson J, et al. Abnormal asymmetry in language association cortex in autism. Annals of Neurology: Official Journal of the American Neurological Association and the Child Neurology Society. 2002; 52(5):588–596.

Hirst RJ, Cragg L, Allen HA. Vision dominates audition in adults but not children: A meta-analysis of the Colavita effect. Neurosci Biobehav Rev. 2018 Nov; 94:286–301.

Hollestein V, Poelmans G, Forde NJ, Beckmann CF, Ecker C, Mann C, Schäfer T, Moessnang C, Baumeister S, Banaschewski T, Bourgeron T, Loth E, Dell’Acqua F, Murphy DGM, Puts NA, Tillmann J, Charman T, Jones EJH, Mason L, Ambrosino S, et al. Excitatory/inhibitory imbalance in autism: the role of glutamate and GABA gene-sets in symptoms and cortical brain structure. Transl Psychiatry. 2023 Jan; 13(1):18.

Huth AG, de Heer WA, Griffiths TL, Theunissen FE, Gallant JL. Natural speech reveals the semantic maps that tile human cerebral cortex. Nature. 2016 Apr; 532(7600):453–458.

Huth AG, Nishimoto S, Vu AT, Gallant JL. A continuous semantic space describes the representation of thousands of object and action categories across the human brain. Neuron. 2012 Dec; 76(6):1210–1224.

Igelström KM, Webb TW, Graziano MS. Functional connectivity between the temporoparietal cortex and cerebellum in autism spectrum disorder. Cerebral Cortex. 2017; 27(4):2617–2627.

Jolly E. Pymer4: Connecting R and python for linear mixed modeling. J Open Source Softw. 2018 Nov; 3(31):862.

Kember J, Patenaude P, Sweatman H, Van Schaik L, Tabuenca Z, Chai XJ. Specialization of anterior and posterior hippocampal functional connectivity differs in autism. Autism Res. 2024 May; .

Keown CL, Shih P, Nair A, Peterson N, Mulvey ME, Müller RA. Local functional overconnectivity in posterior brain regions is associated with symptom severity in autism spectrum disorders. Cell reports. 2013; 5(3):567–572.

Khosla M, Ngo GH, Jamison K, Kuceyeski A, Sabuncu MR. Cortical response to naturalistic stimuli is largely predictable with deep neural networks. Sci Adv. 2021 May; 7(22).

Klin A, Jones W, Schultz R, Volkmar F, Cohen D. Visual fixation patterns during viewing of naturalistic social situations as predictors of social competence in individuals with autism. Archives of general psychiatry. 2002; 59(9):809–816.

Koldewyn K, Whitney D, Rivera SM. Neural correlates of coherent and biological motion perception in autism. Dev Sci. 2011 Sep; 14(5):1075–1088.

Kriegeskorte N, Douglas PK. Interpreting encoding and decoding models. Curr Opin Neurobiol. 2019 Apr; 55:167–179.

Lahnakoski JM, Glerean E, Salmi J, Jääskeläinen IP, Sams M, Hari R, Nummenmaa L. Naturalistic FMRI mapping reveals superior temporal sulcus as the hub for the distributed brain network for social perception. Front Hum Neurosci. 2012 Aug; 6:233.

Lake EMR, Finn ES, Noble SM, Vanderwal T, Shen X, Rosenberg MD, Spann MN, Chun MM, Scheinost D, Constable RT. The Functional Brain Organization of an Individual Allows Prediction of Measures of Social Abilities Transdiagnostically in Autism and Attention-Deficit/Hyperactivity Disorder. Biol Psychiatry. 2019 Aug; 86(4):315–326.

Lawrence KE, Hernandez LM, Bookheimer SY, Dapretto M. Atypical longitudinal development of functional connectivity in adolescents with autism spectrum disorder. Autism Research. 2019; 12(1):53–65.

Lawson RP, Mathys C, Rees G. Adults with autism overestimate the volatility of the sensory environment. Nature neuroscience. 2017; 20(9):1293–1299.

Lawson RP, Rees G, Friston KJ. An aberrant precision account of autism. Front Hum Neurosci. 2014 May; 8:302.

Leekam SR, Nieto C, Libby SJ, Wing L, Gould J. Describing the sensory abnormalities of children and adults with autism. J Autism Dev Disord. 2007 May; 37(5):894–910.

Lefebvre A, Tillmann J, Cliquet F, Amsellem F, Maruani A, Leblond C, Beggiato A, Germanaud D, Amestoy A, Ly-Le Moal M, et al. Tackling hypo and hyper sensory processing heterogeneity in autism: From clinical stratification to genetic pathways. Autism Research. 2023; 16(2):364–378.

Levit-Binnun N, Davidovitch M, Golland Y. Sensory and motor secondary symptoms as indicators of brain vulnerability. Journal of Neurodevelopmental Disorders. 2013; 5(1):26.

Lin F, Albantakis L, Noppari T, Santavirta S, Brandi ML, Sun L, Lukkarinen L, Tani P, Salmi J, Nummenmaa L, et al. Reduced inter-subject functional connectivity during movies in autism: Replicability across cross-national fMRI datasets. bioRxiv. 2025; p. 2025–02.

Lin R, Naselaris T, Kay K, Wehbe L. Stacked regressions and structured variance partitioning for interpretable brain maps. Neuroimage. 2024 Sep; 298(120772):120772.

Lyons KM, Stevenson RA, Owen AM, Stojanoski B. Examining the relationship between measures of autistic traits and neural synchrony during movies in children with and without autism. Neuroimage Clin. 2020 Oct; 28:102477.

Marco EJ, Hinkley LBN, Hill SS, Nagarajan SS. Sensory processing in autism: a review of neurophysiologic findings. Pediatr Res. 2011 May; 69(5 Pt 2):48R–54R.

Martínez K, Martínez-García M, Marcos-Vidal L, Janssen J, Castellanos FX, Pretus C, Villarroya O, Pina-Camacho L, Díaz-Caneja CM, Parellada M, Arango C, Desco M, Sepulcre J, Carmona S. Sensory-to-cognitive systems integration is associated with clinical severity in autism spectrum disorder. J Am Acad Child Adolesc Psychiatry. 2020 Mar; 59(3):422–433.

McClure P, Rho N, Lee JA, Kaczmarzyk JR, Zheng CY, Ghosh SS, Nielson DM, Thomas AG, Bandettini P, Pereira F. Knowing what you know in brain segmentation using Bayesian deep neural networks. Front Neuroinform. 2019 Oct; 13:67.

McDermott J, pycochleagram: Generate cochleagrams natively in Python.; 2018. Accessed: 2024-10-30. https://github.com/mcdermottLab/pycochleagram?tab=readme-ov-file.

Mehta K, Salo T, Madison TJ, Adebimpe A, Bassett DS, Bertolero M, Cieslak M, Covitz S, Houghton A, Keller AS, Lundquist JT, Luo A, Miranda-Dominguez O, Nelson SM, Shafiei G, Shanmugan S, Shinohara RT, Smyser CD, Sydnor VJ, Weldon KB, et al. XCP-D: A robust pipeline for the post-processing of fMRI data. Imaging Neuroscience. 2024 Aug; 2:1–26.

Mihailov A, Philippe C, Gloaguen A, Grigis A, Laidi C, Piguet C, Houenou J, Frouin V. Cortical signatures in behaviorally clustered autistic traits subgroups: a population-based study. Transl Psychiatry. 2020 Jun; 10(1):207.

Moro SS, Ghemraoui AA, Steeves JKE. No Colavita effect: Lack of visual dominance in people with autism spectrum disorder. J Vis. 2012 Aug; 12(9):1038–1038.

Mottron L, Burack JA. Enhanced perceptual functioning in the development of autism. In: Burack JA, Charman T, Yirmiya N, Zelazo PR, editors. The Development of Autism: Perspectives from Theory and Research Lawrence Erlbaum Associates Publishers; 2001.p. 131–148.

Mottron L, Dawson M, Soulières I, Hubert B, Burack J. Enhanced perceptual functioning in autism: an update, and eight principles of autistic perception. J Autism Dev Disord. 2006 Jan; 36(1):27–43.

Mumford D. On the computational architecture of the neocortex: II The role of cortico-cortical loops. Biological cybernetics. 1992; 66(3):241–251.

Napolitano A, Schiavi S, La Rosa P, Rossi-Espagnet MC, Petrillo S, Bottino F, Tagliente E, Longo D, Lupi E, Casula L, et al. Sex differences in autism spectrum disorder: diagnostic, neurobiological, and behavioral features. Frontiers in psychiatry. 2022; 13:889636.

Naselaris T, Kay KN, Nishimoto S, Gallant JL. Encoding and decoding in fMRI. Neuroimage. 2011 May; 56(2):400– 410.

Nastase SA, Goldstein A, Hasson U. Keep it real: rethinking the primacy of experimental control in cognitive neuroscience. Neuroimage. 2020 Nov; 222:117254.

Nebel MB, Lidstone DE, Wang L, Benkeser D, Mostofsky SH, Risk BB. Accounting for motion in resting-state fMRI: What part of the spectrum are we characterizing in autism spectrum disorder? Neuroimage. 2022 Aug; 257(119296):119296.

Neil PA, Chee-Ruiter C, Scheier C, Lewkowicz DJ, Shimojo S. Development of multisensory spatial integration and perception in humans. Developmental science. 2006; 9(5):454–464.

Nieto Del Rincón PL. Autism: alterations in auditory perception. Rev Neurosci. 2008; 19(1):61–78.

Noel JP, Angelaki DE. A theory of autism bridging across levels of description. Trends in Cognitive Sciences. 2023; 27(7):631–641.

Nunez-Elizalde A, la Tour TD, di Oleggio Castello MV, gallantlab/pymoten: v0.0.4. Zenodo; 2022.

O’connor N, Hermelin B. Sensory dominance in autistic imbecile children and controls. Arch Gen Psychiatry. 1965 Jan; 12(1):99–103.

Okada K, Venezia JH, Matchin W, Saberi K, Hickok G. An fMRI Study of Audiovisual Speech Perception Reveals Multisensory Interactions in Auditory Cortex. PLoS One. 2013 Jun; 8(6):e68959.

O’riordan MA. Superior visual search in adults with autism. Autism. 2004; 8(3):229–248.

Pantelis PC, Byrge L, Tyszka JM, Adolphs R, Kennedy DP. A specific hypoactivation of right temporo-parietal junction/posterior superior temporal sulcus in response to socially awkward situations in autism. Soc Cogn Affect Neurosci. 2015 Oct; 10(10):1348–1356.

Pellicano E, Burr D. When the world becomes ‘too real’: a Bayesian explanation of autistic perception. Trends Cogn Sci. 2012 Oct; 16(10):504–510.

Pierce K, Haist F, Sedaghat F, Courchesne E. The brain response to personally familiar faces in autism: findings of fusiform activity and beyond. Brain. 2004; 127(12):2703–2716.

Pierce K, Müller RA, Ambrose J, Allen G, Courchesne E. Face processing occurs outside the fusiformface area’in autism: evidence from functional MRI. Brain. 2001; 124(10):2059–2073.

Puts NA, Wodka EL, Harris AD, Crocetti D, Tommerdahl M, Mostofsky SH, Edden RA. Reduced GABA and altered somatosensory function in children with autism spectrum disorder. Autism Research. 2017; 10(4):608–619.

Rao RP, Ballard DH. Predictive coding in the visual cortex: a functional interpretation of some extra-classical receptive-field effects. Nature neuroscience. 1999; 2(1):79–87.

Redcay E, Moraczewski D. Social cognition in context: A naturalistic imaging approach. Neuroimage. 2020 Aug; 216:116392.

Robertson CE, Baron-Cohen S. Sensory perception in autism. Nat Rev Neurosci. 2017 Nov; 18(11):671–684.

Robertson CE, Ratai EM, Kanwisher N. Reduced GABAergic action in the autistic brain. Current Biology. 2016; 26(1):80–85.

Robertson CE, Thomas C, Kravitz DJ, Wallace GL, Baron-Cohen S, Martin A, Baker CI. Global motion perception deficits in autism are reflected as early as primary visual cortex. Brain. 2014 Sep; 137(Pt 9):2588–2599.

Ronconi L, Vitale A, Federici A, Mazzoni N, Battaglini L, Molteni M, Casartelli L. Neural dynamics driving audiovisual integration in autism. Cereb Cortex. 2023 Jan; 33(3):543–556.

Ross P, Atkins B, Allison L, Simpson H, Duffell C, Williams M, Ermolina O. Children cannot ignore what they hear: Incongruent emotional information leads to an auditory dominance in children. J Exp Child Psychol. 2021 Apr; 204(105068):105068.

Rubenstein JL, Merzenich MM. Model of autism: increased ratio of excitation/inhibition in key neural systems. Genes, Brain and Behavior. 2003; 2(5):255–267.

Rudie JD, Dapretto M. Convergent evidence of brain overconnectivity in children with autism? Cell reports. 2013; 5(3):565–566.

Russo N, Cascio CJ, Baranek GT, Woynaroski TG, Williams ZJ, Green SA, Schaaf R. A cascading effects model of early sensory development in autism. Psychological Review. 2025; .

Rutter M, Bailey A, Lord C. SCQ: The Social Communication Questionnaire. Western psychological services; 2003.

Salmi J, Roine U, Glerean E, Lahnakoski J, Nieminen-von Wendt T, Tani P, Leppämäki S, Nummenmaa L, Jääskeläinen IP, Carlson S, Rintahaka P, Sams M. The brains of high functioning autistic individuals do not synchronize with those of others. Neuroimage Clin. 2013 Oct; 3:489–497.

Samara A, Zada Z, Vanderwal T, Hasson U, Nastase SA. Cortical language areas are coupled via a soft hierarchy of model-based linguistic features. bioRxiv. 2025; p. 2025–06.

Schneebeli M, Haker H, Rüesch A, Zahnd N, Marino S. Disentangling “Bayesian brain” theories of autism spectrum disorder. medRxiv. 2022; .

Schultz RT, Grelotti DJ, Klin A, Kleinman J, Van der Gaag C, Marois R, Skudlarski P. The role of the fusiform face area in social cognition: implications for the pathobiology of autism. Philosophical Transactions of the Royal Society of London Series B: Biological Sciences. 2003; 358(1430):415–427.

Schwarzkopf DS, Anderson EJ, de Haas B, White SJ, Rees G. Larger extrastriate population receptive fields in autism spectrum disorders. J Neurosci. 2014 Feb; 34(7):2713–2724.

Shih P, Keehn B, Oram JK, Leyden KM, Keown CL, Müller RA. Functional differentiation of posterior superior temporal sulcus in autism: a functional connectivity magnetic resonance imaging study. Biological psychiatry. 2011; 70(3):270–277.

Simonoff E, Kent R, Stringer D, Lord C, Briskman J, Lukito S, Pickles A, Charman T, Baird G. Trajectories in symptoms of autism and cognitive ability in autism from childhood to adult life: Findings from a longitudinal epidemiological cohort. Journal of the American Academy of Child & Adolescent Psychiatry. 2020; 59(12):1342–1352.

Smith LE, Maenner MJ, Seltzer MM. Developmental trajectories in adolescents and adults with autism: The case of daily living skills. Journal of the American Academy of Child & Adolescent Psychiatry. 2012; 51(6):622–631.

Sokolov AA, Erb M, Gharabaghi A, Grodd W, Tatagiba MS, Pavlova MA. Biological motion processing: the left cerebellum communicates with the right superior temporal sulcus. Neuroimage. 2012; 59(3):2824–2830.

Sonkusare S, Breakspear M, Guo C. Naturalistic Stimuli in Neuroscience: Critically Acclaimed. Trends Cogn Sci. 2019 Aug; 23(8):699–714.

Stein BE, Stanford TR. Multisensory integration: current issues from the perspective of the single neuron. Nature reviews neuroscience. 2008; 9(4):255–266.

Steinmetz CJ, pyloudnorm: Flexible audio loudness meter in Python with implementation of ITU-R BS.1770-4 loudness algorithm; 2023.

Stevenson RA, Segers M, Ferber S, Barense MD, Wallace MT. The impact of multisensory integration deficits on speech perception in children with autism spectrum disorders. Front Psychol. 2014 May; 5:379.

Stevenson RA, Siemann JK, Woynaroski TG, Schneider BC, Eberly HE, Camarata SM, Wallace MT. Evidence for diminished multisensory integration in autism spectrum disorders. J Autism Dev Disord. 2014 Dec; 44(12):3161–3167.

Supekar K, Uddin LQ, Khouzam A, Phillips J, Gaillard WD, Kenworthy LE, Yerys BE, Vaidya CJ, Menon V. Brain hyperconnectivity in children with autism and its links to social deficits. Cell Rep. 2013 Nov; 5(3):738–747.

Sydnor VJ, Larsen B, Bassett DS, Alexander-Bloch A, Fair DA, Liston C, Mackey AP, Milham MP, Pines A, Roalf DR, et al. Neurodevelopment of the association cortices: Patterns, mechanisms, and implications for psychopathology. Neuron. 2021; 109(18):2820–2846.

Takesian AE, Hensch TK. Balancing plasticity/stability across brain development. Progress in brain research. 2013; 207:3–34.

Tavassoli T, Miller LJ, Schoen SA, Nielsen DM, Baron-Cohen S. Sensory over-responsivity in adults with autism spectrum conditions. Autism. 2014 May; 18(4):428–432.

Thompson RA, Nelson CA. Developmental science and the media: Early brain development. American Psychologist. 2001; 56(1):5.

Tong X, Xie H, Fonzo GA, Zhao K, Satterthwaite TD, Carlisle NB, Zhang Y. Symptom dimensions of resting-state electroencephalographic functional connectivity in autism. Nature Mental Health. 2024 Jan; p. 1–12.

Dupré la Tour T, Eickenberg M, Nunez-Elizalde AO, Gallant JL. Feature-space selection with banded ridge regression. Neuroimage. 2022 Dec; 264:119728.

Dupré la Tour T, Visconti di Oleggio Castello M, Gallant JL. The Voxelwise Encoding Model framework: A tutorial introduction to fitting encoding models to fMRI data. Imaging Neuroscience. 2025; 10.1162/imag_a_00575, doi: 10.1162/imag_a_00575.

Uljarević M, Baranek G, Vivanti G, Hedley D, Hudry K, Lane A. Heterogeneity of sensory features in autism spectrum disorder: Challenges and perspectives for future research. Autism Research. 2017; 10(5):703–710.

Vanderwal T, Eilbott J, Castellanos FX. Movies in the magnet: Naturalistic paradigms in developmental functional neuroimaging. Dev Cogn Neurosci. 2019 Apr; 36:100600.

Wang X, Bouton S, Kojovic N, Giraud AL, Schaer M. Atypical audio-visual neural synchrony and speech processing in children with autism spectrum disorder. bioRxiv. 2024 Apr; p. 2024.04.19.590044.

Wei H, Zhu Y, Wang T, Zhang X, Zhang K, Zhang Z. Genetic risk factors for autism-spectrum disorders: A systematic review based on systematic reviews and meta-analysis. Journal of Neural Transmission. 2021; 128(6):717–734.

Williams ZJ, Schaaf R, Ausderau KK, Baranek GT, Barrett DJ, Cascio CJ, Dumont RL, Eyoh EE, Failla MD, Feldman JI, Foss-Feig JH, Green HL, Green SA, He JL, Kaplan-Kahn EA, Keçeli-Kaysılı B, MacLennan K, Mailloux Z, Marco EJ, Mash LE, et al. Examining the latent structure and correlates of sensory reactivity in autism: a multi-site integrative data analysis by the autism sensory research consortium. Mol Autism. 2023 Aug; 14(1):31.

Xiao Y, Friederici AD, Margulies DS, Brauer J. Longitudinal changes in resting-state fMRI from age 5 to age 6 years covary with language development. Neuroimage. 2016; 128:116–124.

You W, Li Q, Chen L, He N, Li Y, Long F, Wang Y, Chen Y, McNamara RK, Sweeney JA, et al. Common and distinct cortical thickness alterations in youth with autism spectrum disorder and attention-deficit/hyperactivity disorder. BMC medicine. 2024; 22(1):92.

